# Novel substrate prediction for the TAM family of RTKs using phosphoproteomics and structure-based modeling

**DOI:** 10.1101/2023.09.01.555943

**Authors:** Naomi E. Widstrom, Grigorii V. Andrianov, Jason L. Heier, Celina Heier, John Karanicolas, Laurie L. Parker

## Abstract

The TAM family of receptor tyrosine kinases is implicated in multiple distinct oncogenic signaling pathways. However, to date there are no FDA-approved small molecule inhibitors for the TAM kinases. Inhibitor design and screening rely on tools to study kinase activity. Our goal was to address this gap by designing a set of synthetic peptide substrates for each of the TAM family members: Tyro3, Axl and Mer. We used an in vitro phosphoproteomics workflow to determine the substrate profile of each TAM kinase and input the identified substrates into our data processing pipeline, KINATEST-ID, producing a position-specific scoring matrix for each target kinase and generating a list of candidate synthetic peptide substrates. We synthesized and characterized a set of those substrate candidates, systematically measuring their initial phosphorylation rate with each TAM kinase by LC-MS. We also used the multimer modeling function of AlphaFold2 to predict peptide-kinase interactions at the active site for each of the novel candidate peptide sequences against each of the TAM family kinases, and observed that remarkably, every sequence for which it predicted a putatively catalytically competent interaction was also demonstrated biochemically to be a substrate for one or more of the TAM kinases. This work shows that kinase substrate design can be achieved using a combination of preference motifs and structural modeling, and it provides the first demonstration of peptide-protein interaction modeling for predicting likelihood of constructive catalytic interactions.

## INTRODUCTION

The TAM family of receptor tyrosine kinases consists of three members: <u>T</u>yro3, <u>A</u>xl, and <u>M</u>er. They are expressed across a wide range of different tissue types, with their function best studied in immune cells.^1, 2^ The TAM receptors are activated by growth-arrest specific gene 6 (Gas6) and Protein S, among other ligands. The main roles of the TAM family kinases are as mediators of immunosuppressive signaling and the phagocytosis of apoptotic cells, termed efferocytosis. Despite the similarities, these kinases have non-redundant roles in efferocytosis and different expression patterns across innate immune cells.^3, 4^

Although the TAM family kinases haven’t traditionally been labeled as oncogenes, an overwhelming amount of evidence suggests that they play a key role in supporting cancer progression. The TAM receptors are overexpressed or ectopically expressed in a large variety of solid and hematological cancers, and their expression has been correlated to different aspects of the varying cancers, such as decreased survival, metastasis, and chemoresistance.^5^ While their specific role often varies depending on the cellular context, the TAM receptors are typically associated with pro-survival signaling, invasion, and migration. Axl has been implicated in metastasis, migration, invasion and drug resistance in a number of different cancers.^6, 7^ Mer activity has also been tied to motility and survival signaling under varying contexts.^8-11^ There are fewer studies on the individual role of Tyro3 compared to Axl and Mer, but despite being understudied, Tyro3 remains an important cancer target as well.^12-16^

The TAM kinases are often treated interchangeably, but studies reveal subtle differences in signaling leading to differential downstream effects. In tumor types where multiple TAM receptors are expressed, they may have both overlapping and non-overlapping roles, suggesting independent mechanisms supporting cancer survival and growth.^17-19^ Additionally, due to the physiological role the TAM family plays in immune suppression, their expression in innate immune cells within the tumor microenvironment (TME) may help cancers grow and evade immune detection.^20^ Therefore, even when not expressed within cancer cells, TAM activity can still aid cancer growth. Preliminary testing of combination therapies suggests dual inhibition of TAM kinases and checkpoint blockade therapeutics may increase efficacy of treatments.^21-24^

Despite these studies, much about the physiological and oncogenic function of the TAM family kinases remains unknown. To aid the study of these kinases and inhibitor screening efforts, our goal was to design synthetic peptide substrates capable of providing a readout of TAM family kinase activity for in vitro kinase assays. Using a previously established phosphoproteomics workflow based on in vitro phosphorylation of a protease digested cell lysate^25-27^ (with slight adaptations as described in the Experimental section), we determined the substrate profile for each TAM family kinase and used our KINATEST-ID V2.1.0 R-package to analyze the amino acid preferences of each kinase. This allowed us to evaluate the in vitro amino acid preferences of the TAM family via position-specific scoring matrices (PSSMs) and design a set of 15 synthetic peptide substrate candidates based on those PSSMs. We analyzed the phosphorylation of the 15 candidate synthetic substrates against each kinase and found some differential preferences between the three kinases that could not fully be explained by their PSSM scoring. Therefore, we also developed an AlphaFold2 multimer modeling approach to assess the likelihood that each candidate to engage in catalytically favorable interactions with each TAM kinase domain. In this work, we describe how this combination of strategies resulted in development of a set of novel peptide substrates for TAM kinases (Figure 1), and remarkably accurate prediction of the ability of Tyro3, Axl and Mer to phosphorylate these sequences.

**Figure 1.**
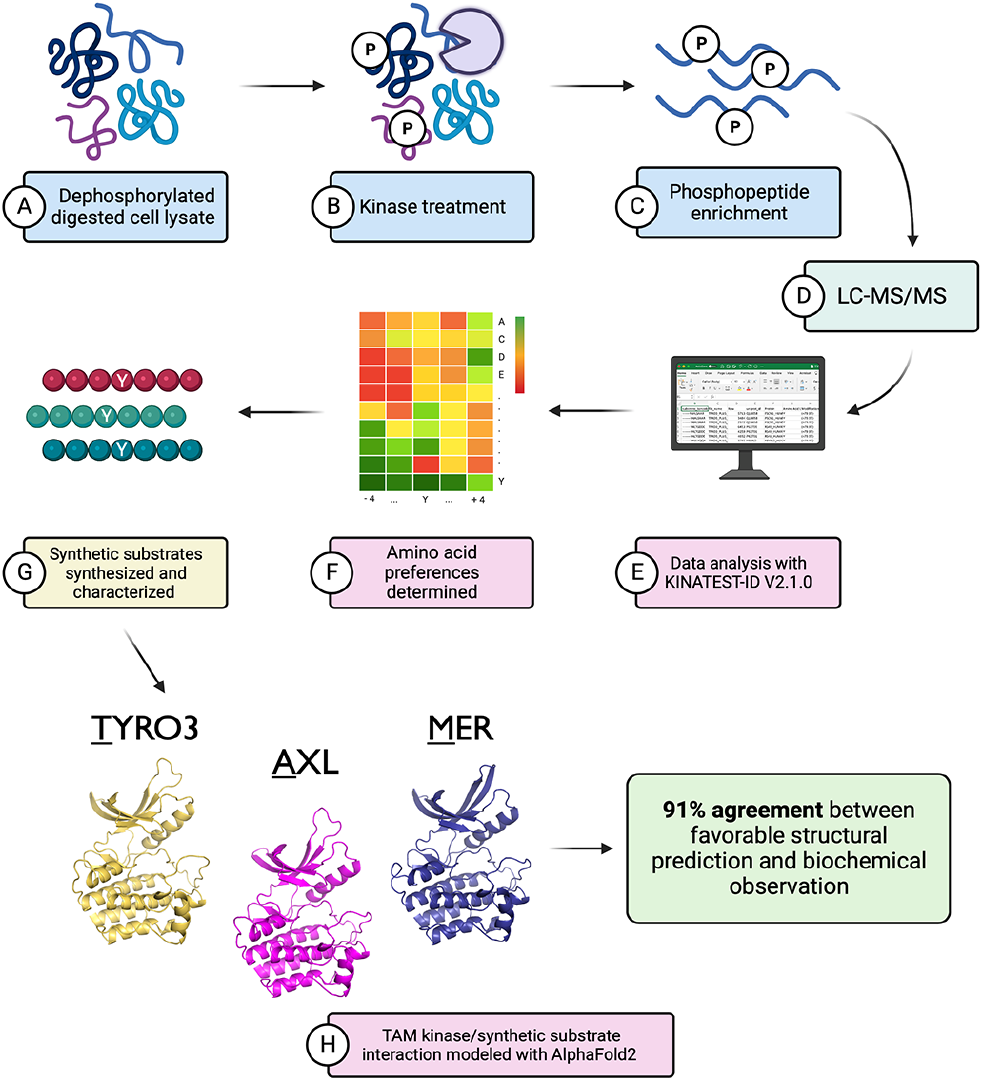
Schematic overview of workflow. A.) Dephosphorylated digested cell lysate used as a natural peptide library treated with B.) recombinant kinase (Tyro3, Axl, or Mer) for 2 hours and C.) subsequent phosphopeptides enriched and D.) identified via LC-MS/MS. E.) Identified peptides processed and input into R-package KINATEST-ID, which F.) determines amino acid preferences used to generate an in-silico library of putative substrates. G.) Synthetic substrates are chosen and synthesized. Created with Bio-Render.com

## RESULTS

### TAM family peptide substrate profiling with in vitro phosphoproteomics

In order to design novel substrates, sufficient information about a kinase’s substrate preference profile needs to be available. Despite the advances made in elucidating the function of TAM family kinases in both physiologically healthy and diseased contexts, the total number of previously reported TAM family substrates remains low, with a total of 26 known sites between all three kinases.^28^ Previously, we have successfully used an adapted in vitro phosphoproteomics workflow to expand the number of known substrates, identify the preferences and design synthetic peptide substrates for target kinase that lacked sufficient numbers of reported substrates.^25, 29^ To determine the substrates for the TAM family kinases, cell lysate was reduced and alkylated, digested with trypsin, and treated with lambda phosphatase to dephosphorylate endogenous phosphosites. This was used as a natural peptide library, reacted in kinase reaction buffer (as described in the Experimental Section) for 2 hrs in triplicate with recombinantly prepared enzyme comprising the intracellular portion of each: Tyro3 (455-end), Axl (473-end), and Mer (528-end). Following phosphopeptide enrichment of the resulting material, samples were analyzed on an Orbitrap Velos mass spectrometer and sequences identified by PEAKS Studio X Pro. The identified phosphopeptides were used as input into our KINATEST-ID V2 pipeline.^25^ In brief, KINATEST-ID V2 creates a Fisher Odds-based PSSM for each amino acid at positions surrounding a central phosphorylation site. To ensure that the most robustly observed peptides were used for these calculations, the input phosphopeptide sequences were filtered to only include those found in all three replicates, and removing any residual endogenous phosphopeptides from the cell lysate tryptic digest observed in the controls that were not treated with kinase.

While the TAM kinases showed some similarities in substrate profiles based on this in vitro phosphorylation experiment, they each had unique substrates identified as well (Figure 2A and Tables S1-S3). Comparison of the PSSMs showed both shared and unshared preferences for certain amino acids (Figure 2B). In particular, relative to the central tyrosine, all three kinases preferred aspartic acid (D) at -3, asparagine (N) at -2, histidine (H) at -1, alanine (A) and glutamic acid (E) at +1, and leucine at +3. On the other hand, each kinase also had preferences for amino acids at some positions that were different than the other family members, which suggested substrates could potentially be designed that could be preferentially phosphorylated by an individual TAM kinase.

**Figure 2.**
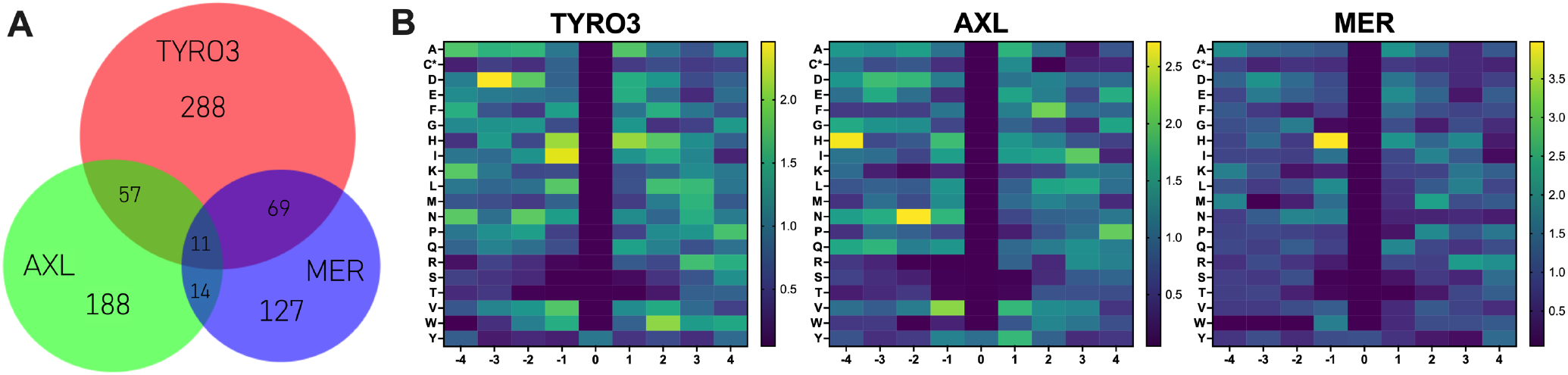
Summary of phosphopeptides identified and heatmaps of Fisher Odds values for Tyro3, Axl, and Mer. A) Venn diagram illustrating shared and unique phosphopeptides identified in all three replicates of in vitro phosphorylation reactions for each kinase (but that were not observed in the input control). B) PSSMs based on Fisher Odds analysis were mapped and color-coded for every amino acid (y-axis) in each position relative to central tyrosine (x-axis). Values above 1 indicate overrepresentation of the amino acid at the position, and values below 1 indicate an underrepresentation. *C = carbamidomethylcysteine.

### Candidate peptide substrate design and characterization

We then used the PSSMs to select amino acids to include for each position, and applied the *Generator* function to permute the motifs and create a list of candidate sequences for each TAM kinase. These were scored against Tyro3, Axl, and Mer’s preferences, and also against the internal panel of tyrosine kinase preferences we have accumulated in KINATEST-ID, using the *Screener* function. We did some manual assessment amongst those options to choose motifs that initially appeared to maximize target kinase preferences, and designed an initial panel of fifteen synthetic substrates in total: six based on the observed Tyro3 preference motif, four on the Axl motif, and five on the Mer motif (Table 1). Not all were designed around the -4 to +4 range that we use for calculating KINATEST-IDV2.1.0 scores (rationale for each set described in sub-sections below). Each substrate was synthesized (as described in the Experimental Section) with biotinylated lysine on the C-terminus, flanked by glycine spacers, to allow affinity capture of the peptides for antiphosphotyrosine antibody-based initial activity screening. The candidate substrates were scored from -4 to +4, and a threshold for likely activity was applied (scores above the threshold indicated with underlining in Table 1). The threshold is based on comparison of Fisher odds-based scores for observed phosphopeptides from the in vitro phosphoproteomics KALIP experiment, vs. tyrosine-containing peptides from the full-length proteins represented that can reasonably be considered likely present, but not observed as phosphopeptides (see the detailed description of this approach in Widstrom et. al.^25^) and graphical representations of the threshold cutpoints in the KINATEST-ID V2.1.0 outputs provided in the supporting files). Based on preliminary KINATEST-ID 2 versions that examined amino acid preferences in different frames around the central tyrosine, the Tyro3 candidates only extend to the -3 position, and some of the substrate candidates have amino acids extended beyond the -4 to +4 frame. In all cases, however the scores reported in Table 1 were ultimately calculated using KINATEST-ID V2.1.0.

**Table 1.**
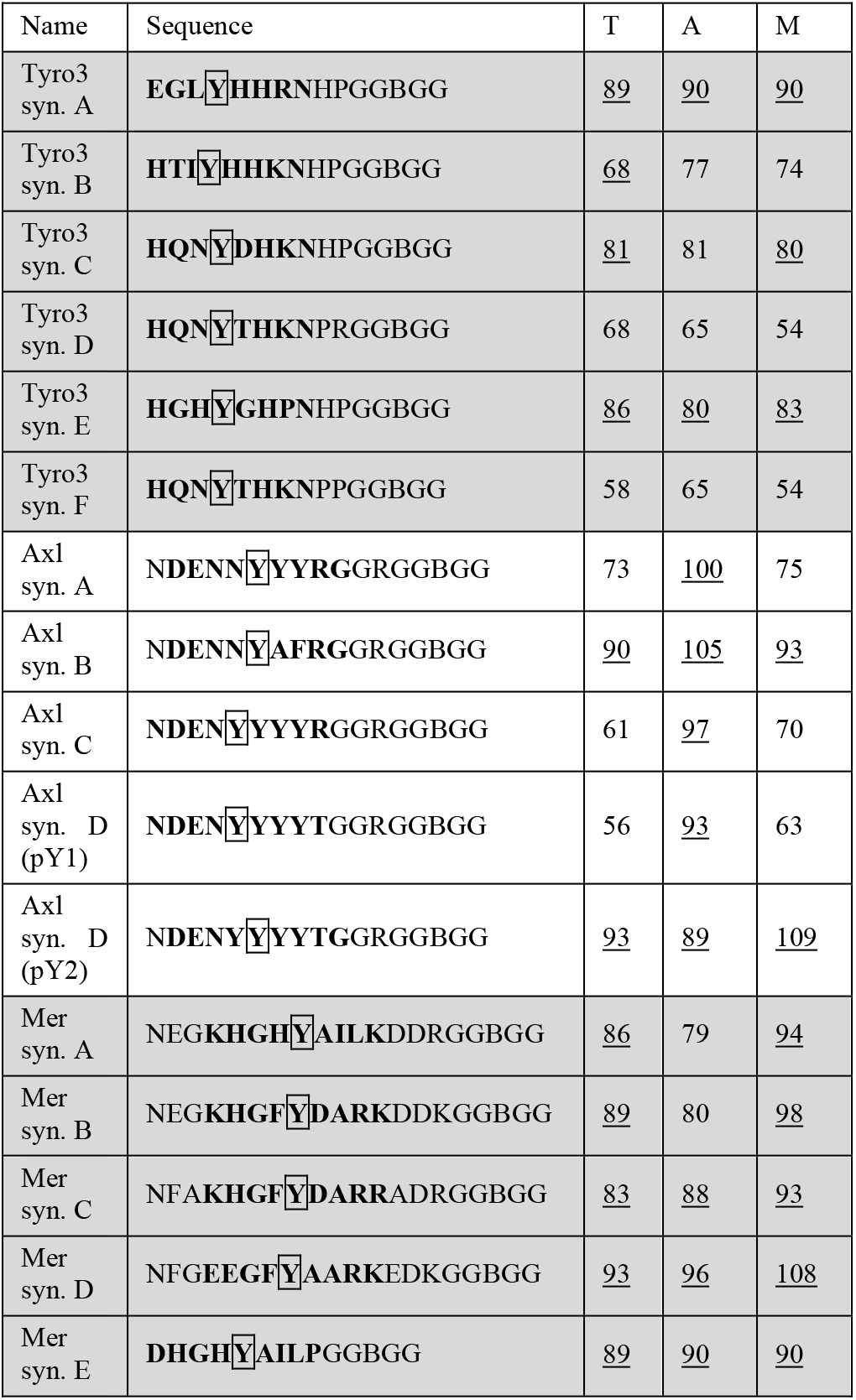
Synthetic substrate (syn.) sequences and scores for major phosphorylation sites. Bold text indicates portion of peptide used to score each substrate. 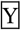 indicates phosphorylation site. Underlined scores indicate score is above threshold for target kinase (see ref.^25^for details). B denotes biotinylated lysine.

#### Tyro3 candidate synthetic substrates

The sequences used in the Tyro3-targeted synthetic substrates (Tyro3 syn. A-F) were chosen with an aim for specificity over efficiency of phosphorylation. We included substrates with lower scores (below KINATEST-ID V2.1.0’s predicted threshold of activity), that had higher predicted specificity (i.e., might favor Tyro3 over the others despite being less efficient substrates). Because we observed many peptides from the Tyro3 reaction that had phosphorylation at a tyrosine within 2-3 amino acids of the N-terminus (Fig. S1), we opted to design the Tyro3 substrates starting at -3 from the phosphosite. To test the putative Tyro3 substrates, we performed an in vitro kinase assay (as described in the Experimental Section) with recombinant Tyro3, quenching aliquots at time points to monitor reaction progress across 60 minutes (Figure 3A). We used an ELISA-based approach, capturing the peptide in Neutravidin-coated plates and detecting phosphorylation with anti-phosphotyrosine antibody 4G10. We found Tyro3 syn. A had the highest rate of phosphorylation and overall signal compared to the other substrates in this experiment. Of the other substrates, by 60 minutes Tyro3 syn. D and E showed moderate signal, B and F low signal, while syn. C remained unphosphorylated.

**Figure 3.**
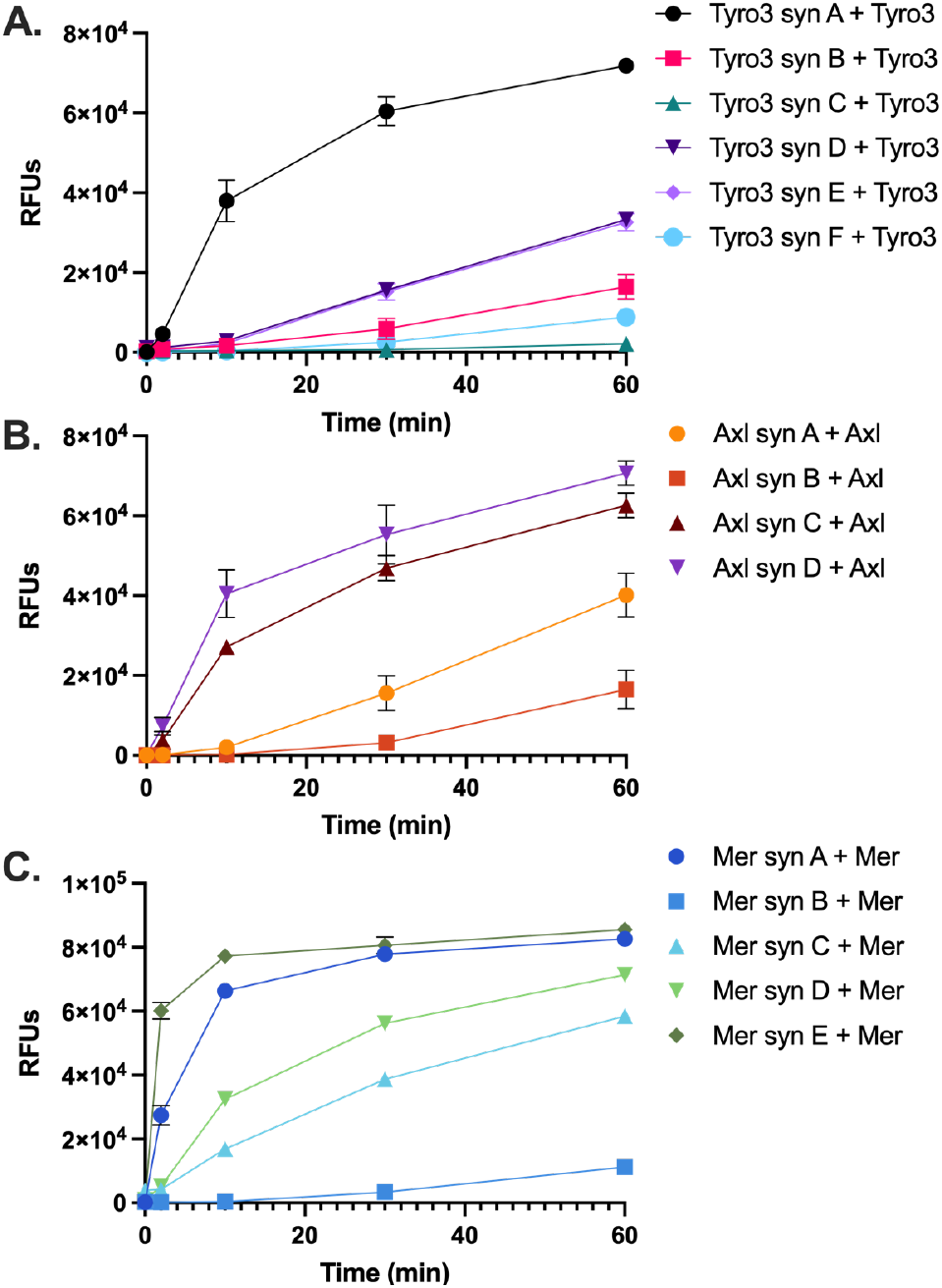
Peptide phosphorylation by TAM family targets and other kinases measured by phosphotyrosine ELISA. Kinase assays were conducted using conditions as described in the Experimental Section, with EDTA-quenched timepoints incubated in 96-well streptavidin-coated plates to capture biotinylated peptide. Phosphorylation analyzed using 4G10 antiphosphotyrosine antibody and Amplex Red as described in the Experimental Section. (A) Tyro3-targeted substrates with recombinant Tyro3. (B) Axl-targeted substrates with recombinant Axl. (C) Mer-targeted substrates with recombinant Mer.

#### Axl candidate synthetic substrates

Compared to either Tyro3 or Mer, the Axl amino acid preferences indicated N in the N-terminal region and additional tyrosine residues near the central phosphosite were preferred. The four Axl synthetic substrates were thus designed to examine a potential preference for multiple tyrosines and tested with an in vitro kinase assay across a 60-minute time course (Figure 3B). The presence of multiple tyrosines adds an additional caveat, as different or multiple sites could be phosphorylated. With the ELISA-based readout method, phosphorylation of different tyrosine sites is indistinguishable, as the anti-pY antibody used will bind to any pY sites present—and due to steric hindrance multiple phosphotyrosines in the epitope would likely still only bind one molecule of antibody per peptide, so differential phosphorylation can’t be assessed from these data. However, the ELISA readout approach was suitable as an initial rapid comparison of peptide candidates, and site-specific phosphorylation was elucidated in the mass spectrometry-based assay described later in this work. Of the substrates, both Axl syn. C and D showed rapid initial phosphorylation signal, while Axl syn. A and B had much lower initial phosphorylation but did show signal by 60 minutes.

#### Mer candidate synthetic substrates

For Mer, four initial peptides were designed for the full span from -7 to +7, based on the KINATEST-ID analyses, while the fifth substrate (Mer syn. E) is a modified shorter (-4 to +4) substrate candidate adjusted to be more similar to universal peptide 5 (U5)^30^, which was observed to be very rapidly phosphorylated by Mer in control experiments (Fig. S2). The in vitro Mer kinase assay shows Mer syn. E and A had very rapid initial phosphorylation, syn. B with very minimal signal after 60 mins, and the rest with moderate to high phosphorylation by Mer (Figure 3C).

### Evaluation of TAM family specificity of the synthetic substrates

Next, to gain insight into the TAM specificity of each substrate and provide further clarity into the individual substrate preferences of each TAM family member, we evaluated off-target phosphorylation by the other TAM kinases for each of the substrate candidates. To do this, we performed one-pot kinase reactions with all 15 synthetic substrate candidates with each of the TAM kinases, either Tyro3, Axl, or Mer (details provided in the Experimental Section). Aliquots from the reaction were quenched at 0, 1, 3, 5, 10, and 20 minutes and the product formation for each peptide was monitored with LC-MS. The extracted ion chromatogram areas (EICs) for each substrate peptide and its corresponding phosphorylated product were quantified, and for each substrate, the phosphopeptide signal was normalized as the ratio of phosphopeptide product signal to total (phosphopeptide plus unphosphorylated) peptide signal and plotted against time (Fig. S3). The resulting slope (normalized EIC/time) was used as a surrogate metric for phosphorylation efficiency. These slopes were plotted in a heatmap (Figure 4). In general, the reactions of each TAM kinase plus their matching substrate candidates (e.g., Tyro3 kinase plus Tyro3 substrates) recapitulate trends from experiments that used streptavidin-based ELISA (Fig. 3).

**Figure 4.**
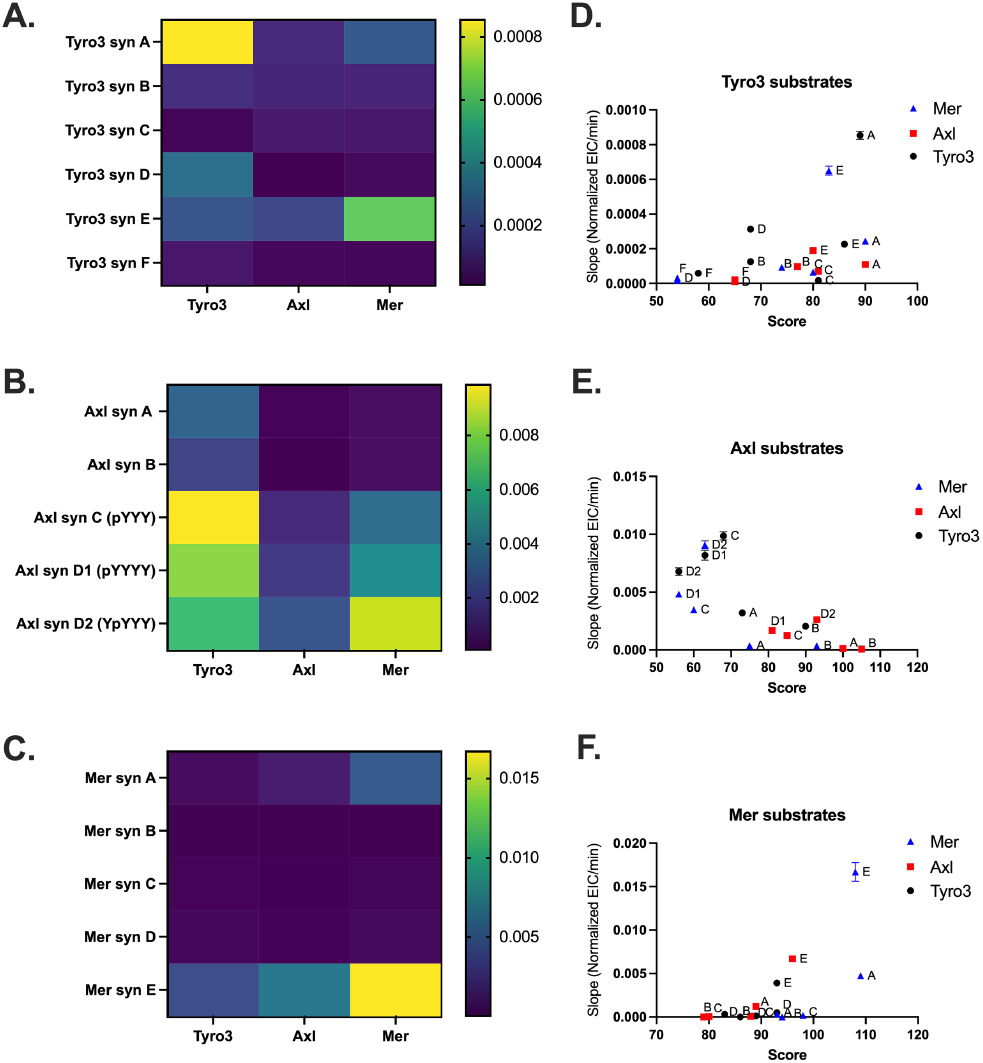
Initial phosphorylation rates of TAM synthetic substrates by Tyro3, Axl, or Mer. In vitro kinase assay with the 15 synthetic substrates in a one-pot reaction with either recombinant Tyro3, Axl, or Mer. Timepoints were collected at 0, 1, 3, 5, 10, and 20 minutes in triplicate and analyzed with LC-MS. Product signal (defined as EIC area) was normalized as product/total (substrate + product) per peptide (Normalized EIC) and plotted against time. Slope values (Normalized EIC/time) were used as metrics for the rate of phosphorylation for each combination of substrate and kinase. Major and minor products of Axl substrates represent different single phosphorylation sites, identified by differences in LCMS retention time. Heatmaps A, B and C: Visualized comparisons for the increase in phosphorylation signal over time (normalized EIC/min) for Tyro3, Axl, or Mer and A.) Tyro3 synthetic substrates, B.) Axl synthetic substrates and C.) Mer synthetic substrates. Graphs D, E and F: Plots of KINATEST-ID V2.1.0 score against increase in phosphorylation signal over time (normalized EIC/min) for reaction of recombinant Tyro3, Axl, and Mer with D.) Tyro3 synthetic substrates, E.) Axl synthetic substrates, and F.) Mer synthetic substrates. Data represented as slope (normalized EIC/min) ± standard error of the mean (SEM).

#### Specificity of Tyro3-targeted sequences

For the Tyro3 substrates, Tyro3 syn. A had the highest rate of phosphorylation by Tyro3, while Tyro3 syn. D and E showed lower levels of phosphorylation, similar to previous results. Off-target reactions show that Mer also phosphorylates Tyro3 syn. A, and more notably phosphorylates Tyro3 syn. E at a higher rate than Tyro3. Axl also phosphorylated Tyro3 syn. E, to a comparable level as the Tyro3 reaction, but the rest of the substrates seemed to be poor substrates for Axl (Figure 4A).

#### Specificity of Axl-targeted sequences

The multiple tyrosine residues in Axl syn. A, C and D added complexity to the analysis as there is the possibility that any of the three (Axl syn. A) or four (Axl syn. C and D) tyrosine residues could be phosphorylated. To establish the modification sites for each of these peptide products, we initially differentiated the isobaric parent ions for the different products by MS/MS fragmentation, and noted that they exhibited different retention times, which later enabled individual analysis without the use of MS/MS. Axl syn. A only has one major product (pYYY), as the other possible products had very low to non-detectable background signal. Axl syn. C has distinguishable major (pYYY) and minor (YpYY) products, while Axl syn. D has two major products (pYYYY and YpYYY), suggesting two different tyrosines were phosphorylated roughly equally. We also observed very minor double phosphorylation of Axl syn. C and D but not triple or quadruple phosphorylation. Overall, we found the Axl substrates were phosphorylated at a higher rate by Tyro3 or Mer compared to Axl (Figure 4B). However, it is possible that under these conditions the Axl enzyme was overall less active than Tyro3 or Mer. Of the Axl substrates, Axl syn. D had the highest rate of phosphorylation by Axl, followed by Axl syn. C. Tyro3 phosphorylated Axl syn. C at the highest rate, and with a higher rate than any of the Tyro3 substrates. Tyro3 also phosphorylated both Axl syn. D products fairly equally. Interestingly, while Mer also phosphorylated both Axl syn. D products, one of the sites (YpYYY) was apparently phosphorylated more efficiently than the other site (major product 1). Overall, none of these substrates appear to be specific for Axl, although Axl syn. A has potential for Tyro3 specificity.

#### Specificity of Mer-targeted sequences

Of the Mer synthetic substrates, Mer syn. E had the highest rate of phosphorylation by all three kinases (Figure 4C). Although Mer syn. E lacks specificity, it still shows a rapid initial rate of phosphorylation-that may be very useful for in vitro kinase assays or TAM family screening efforts. Mer syn. A is also efficiently phosphorylated by Mer, and minorly phosphorylated by Axl. At these timepoints, none of the rest of the substrates demonstrated significant phosphorylation.

Comparing all three sets of substrates, the phosphorylation rate trends by Mer and Axl appear somewhat consistent, although Axl phosphorylation is consistently lower. This may suggest that the Axl enzyme itself was relatively less active under these conditions compared to Mer. Considering the amino acid preferences determined by KINATEST-ID, the Axl substrate sequences contained multiple positions with amino acids favored by all three TAM kinases. This may have contributed to the non-specific nature of these synthetic substrates. In contrast, the Tyro3 synthetic substrate sequences contained fewer pan-TAM favored amino acids, instead including amino acids favored only by Tyro3 at specific positions. This may have helped the specificity of these substrates. However, compared to the other substrates the Tyro3 substrates were all phosphorylated at a lower rate. These substrates were designed with specificity in mind, which in this case may have resulted in a tradeoff with efficiency. Overall, this comparison of 15 different substrates reveals subtle differences in the substrate preferences of the three highly related kinases, and may inform selection of substrates for future development in TAM family member-specific assays.

### Evaluation of the KINATESTID V2.1.0 PSSM capabilities for substrate prediction

Next, we evaluated the correlation between slope of the phosphorylation signal and each sequence’s KINATEST-ID score (Table 1). Both the Tyro3 (Figure 4D) and Mer (Figure 4F) synthetic substrates had a significant positive correlation between substrate score and performance. However, we did not observe a correlation between score and initial slope for the Axl synthetic substrates (Figure 4E). A reason for this may be that the ability of the Fisher odds PSSMs to predict substrate behavior for each of these kinases depends on the composition of its input dataset (Table S4). The Axl PSSM was built from an observed substrate list that contained ∼6% sequences with at least one Y at +1 from the phosphosite, whereas the Tyro3 and Mer PSSMs were built from lists of sequences in which only 1-2% had that pYY characteristic. Given that the Axl-designed peptides all contained multiple sequential Y residues, the Tyro3 and Mer PSSMs were likely not able to predict substrate behavior for those sequence options accurately. In an attempt to develop a more accurate prediction approach, we moved to a structural modeling strategy and assessed how well the phosphorylation of these 15 peptides by each of the TAM family kinases could be predicted.

### Structural prediction of substrates using AlphaFold2

With the advent of AlphaFold2 (AF2), we now have unprecedented access to structural information for protein-protein complexes, in many cases at resolution comparable to experimentally-determined structures.^31, 32^ Inspired by this development, we evaluated whether the current version of AF2 could accurately predict kinase-substrate interactions for our new panel of peptide sequences.

At the outset, we noted that AF2 consistently predicts inactive conformations for certain kinases, irrespective of which substrate peptides they are modeled with (and at times in the absence of any peptide). AF2 models can be steered towards specific conformations of a given target by tuning the multiple sequence alignment (MSA) that it uses as input, adjusting structural templates, or both. When building peptide-bound models of each kinase, we sought to generate maximally unbiased predictions from AF2, by electing to use single sequence inputs (i.e. no MSA data) but providing a catalytically active template for each kinase. Accordingly, we began by using AF2 with no substrate present, and using MSA data alone to generate models of Tyro3, Axl, and Mer in their active conformations. For each kinase, we then used AF2 to build models of each kinase with each synthetic substrate, using this kinase’s AF2-generated active conformation as a template.

To allow for direct comparisons between experimentally-derived catalytic efficiency and computationally-predicted structural models, we began by converting both to binary representations. For catalytic efficiency, we classified binders versus non-binders using a threshold value for the rate of 0.0002 product normalized EIC/min (from the pooled substrate assay described above, Figure 4), leading to a map of which synthetic peptides could serve as a substrate for each kinase (Figure 5A). For the AF2 models, we classified each model on the basis of whether the output conformation was suitably oriented for catalysis (Figure 5B): models in which the peptide was not correctly placed for catalysis, or on which the kinase was not in an active conformation, were interpreted as predictions that this peptide/kinase pairing would not be catalytically active. Specifically, interpreting a given model as”predicted active” required that the Tyr from the peptide be placed close to the Asp of the kinase’s HRD motif (a structural feature known to be required for phosphate transfer onto the Tyr), and that the kinase adopts precisely the conformation needed for binding ATP, magnesium, and substrate ^33^ (Figure 5C).

**Figure 5.**
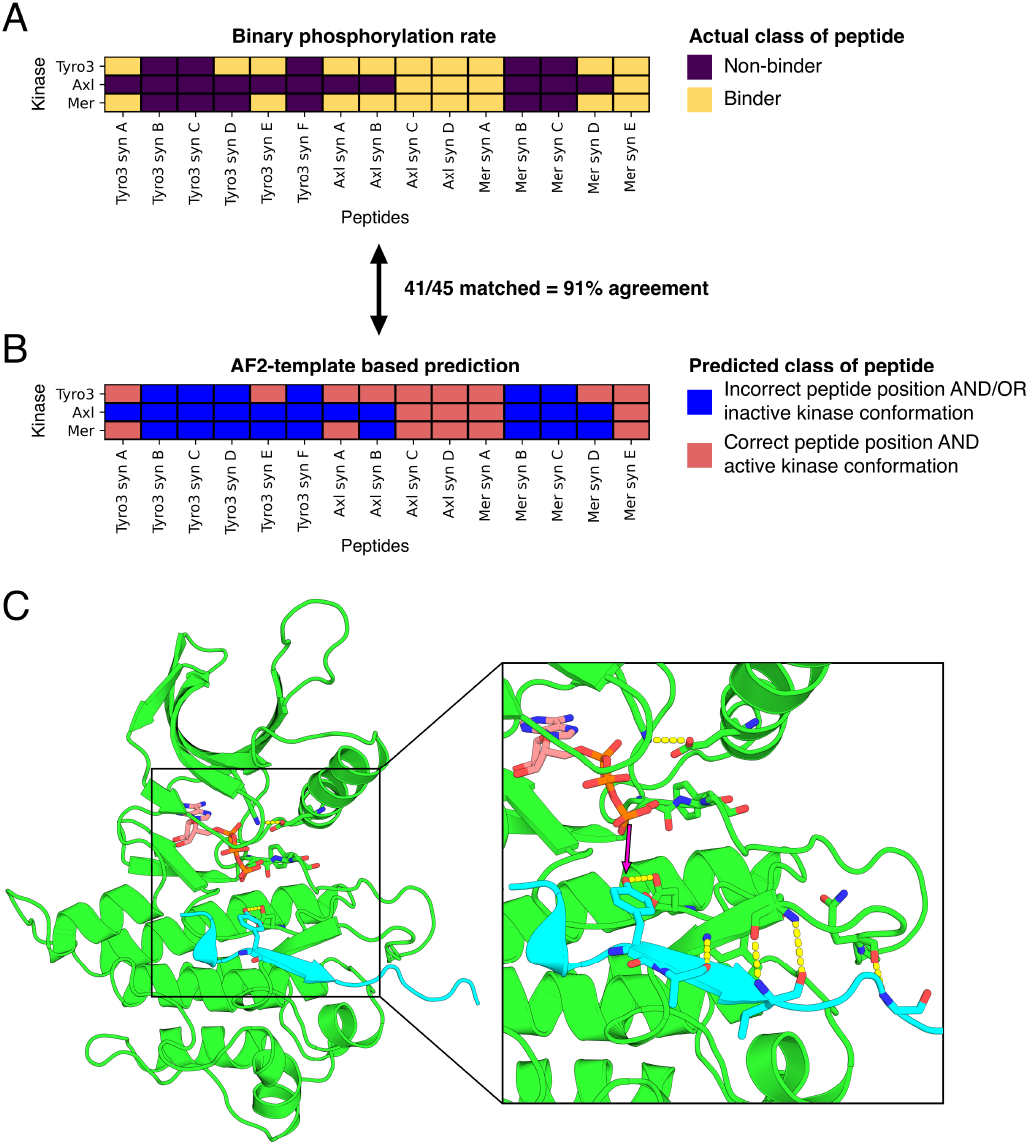
AlphaFold2 modeling to predict catalytically productive kinase-substrate complexes. A) Transformation of phosphorylation rates into discrete classes (non-binders versus binders). B) Predicted classes of kinase-peptide pairs by AlphaFold2 using 3D templates. C) Representative model for one of the pairings predicted to be catalytically active, the Mer syn E peptide (cyan) bound to the active conformation of Tyro3 (green).

Strikingly, the accuracy of the AF2 predictions was 91% (Figure 5), with no false positive predictions. Only four peptides were incrorrectly predicted as non-binders: for the Tyro3 kinase our AF2 models did not correctly identify Tyro syn. D as a substrate, and for Mer kinase our models did not find Tyro syn E, Axl syn. B or Mer syn. C (Figure 5). Interestingly, the AF2 models did occasionally predict phosphorylation of residues other than the canonical”central” -4 to +4 tyrosine in the Tyr-rich sequences, such as Axl syn. C and D, which was also seen in the biochemical analyses for those Axl-targeted substrates for which more than one product was observed.

Overall, these AF2 models exhibit remarkable recapitulation of even the subtle details from experimentally-derived kinase substrate preferences. For example, the Tyro3 syn. A is correctly recognized as a substrate for the Axl kinase alone; meanwhile, Tyro syn. B is correctly recognized as a substrate for all three kinases (Figure 5). Collectively, this concordance of the experimental observations could only arise from correctly predicting structural details of the extensive contacts between the kinase and its substrates. To our knowledge, this is the first demonstration of using structure-based modeling to predict kinase substrate pairings, and our method achieves this task with unprecedented accuracy.

## DISCUSSION

The TAM family of kinases are implicated in several different cancers, typically for their roles in migration, metastasis, and chemoresistance. As regulators of immune suppression, over-activation of the TAM receptors may contribute to an immuno-suppressive tumor microenvironment, which aids tumor growth and progression. Their activity may also prevent infiltrating immune cells from initiating immune destruction of tumor cells.^20^ TAM inhibitors therefore may be effective at both treating tumor progression and in combination with other therapeutics to increase efficacy of treatments. However, to date there are no FDA-approved TAM kinase small molecule inhibitors. Additionally, despite the interest in the TAM family for cancer related activity, few substrates of each kinase are known, adding difficulty to studying the activity of the kinases for in vitro kinase assays or inhibitor screening assays.

### Synthetic substrate peptides for the TAM family of kinases

Overall, we successfully designed a set of synthetic peptide substrates that can be used to probe the activity of the TAM family kinases using in vitro kinase assays. We further delineated the differences between Tyro3, Axl, and Mer activities by demonstrating distinct amino acid preferences and by providing direct comparison of synthetic substrate preferences. From the pooled substrate LC/MS assay, we observed several substrates that may be good candidates for differentiating between TAM family members. For example, Tyro3 syn. A was rapidly phosphorylated by Tyro3, with minor signal from Axl and Mer by 30 minutes. While Tyro3 syn. D had lower signal overall by 30 minutes, it showed no phosphorylation by Axl or Mer. Despite the lower efficiency of phosphorylation, Tyro3 syn. D may be a good candidate for selectively detecting Tyro3 activity in the presence of the other TAM family members. This provides a set of tools to further study the TAM kinases in vitro, which could be applied towards inhibitor screening efforts, and presents evidence for subtle differences in the TAM family kinase substrate preferences. One difficulty in applying these in more complex assays is the lack of known substrates for other kinases to compare the putative sequences against. As more substrates are identified for a larger range of kinases, it may become easier to select sequences more likely to be specific. However, due to the conserved nature of kinases, especially between family members, total substrate specificity may not be achievable for every kinase. On top of substrate binding, the active site of a kinase must allow for coordination of ATP and Mg^2+^, and active orientation of the activation loop and αC helix. As a result, kinase active sites are very similar, making it difficult to differentiate on the basis of phosphorylation site specificity alone. Recent work by Johnson et al. demonstrated that there are”negative selection” factors at some positions for many serine/threonine kinases^34^—this may also be possible to exploit for achieving selectivity for tyrosine kinases in future substrate design applications.

### Structural modeling as an approach for candidate substrate evaluation

The development of AlphaFold2 revolutionized the field of structural biology. By providing accurate structural predictions of the kinase-peptide complex, AlphaFold2 can facilitate the selection of promising peptide candidates for experimental validation. Moreover, the structural insights gained from AlphaFold2 predictions can also guide the design of improved peptides with higher affinity and specificity for their target kinases.

## EXPERIMENTAL SECTION

### Cell culture and phosphoproteomics sample preparation

K562 (ATCC) cells were grown and maintained in IMDM (Gibco) supplemented with 10% fetal bovine serum (FBS), 1% penicillin/streptomycin in 5% CO_2_ at 37°C. Cells were washed in phosphate buffered saline (PBS) and resuspended in lysis buffer (7M urea, 2M thiourea, 0.4 M Tris pH 7.5, 20% acetonitrile, 4 mM tris(2-carboxyethyl)phosphine (TCEP), 5 mM ethylenediaminetetraacetic acid (EDTA), 10 mM sodium fluoride, 1 mM solidum orthovanadate, 1X Halt Phosphatase inhibitor cocktail (ThermoFisher)). Lysed cells were probe sonicated and alkylated by adding chloroacetamide (Sigma-Aldrich) to a final concentration of 8mM. Samples were centrifuged at 15,000 x g at 4 °C for 15 minutes to remove insoluble cellular debris. For lysate digestion, samples were diluted 5-fold by adding 25 mM Tris, pH 8.0 and trypsin protease (ThermoFisher) added in a 1:40 ratio and incubated overnight at 37°C. Digestion was quenched by adding an equal volume of 0.6% trifluoroacetic acid (TFA) to solution and samples desalted with Oasis HLB 1 cc Vac Cartridge (Waters). To remove endogenous phosphate groups, samples were reconstituted in phosphatase reaction buffer (50 mM 4-(2-hydroxyethyl)-1-piperazineethanesulfonic acid (HEPES), 100 mM NaCl, 2 mM Dithiothreitol (DTT), 1 mM MnCl_2,_ pH 7.5) and 4000 U lambda protein phosphatase (New England Biolabs) added per 1 mg digested lysate. Samples incubated at 30°C overnight and deactivated by heating at 65°C for one hour. Each sample was equally split in two, one half to be used as the (+) kinase treatment and the other as the (-) kinase control. Activity of each kinase was validated beforehand using universal tyrosine kinase substrate U5^30^ via the methods described in the”In vitro synthetic substrate kinase assay with ELISA readout” paragraph (Fig. S2). Kinase reaction buffer added to each sample (final concentrations of 50 mM Tris pH 7.5, 10 mM MgCl_2_, 1 mM DTT, 1 mM Na_3_VO_4_, 2 mM ATP) and 2 ug of recombinant kinase (SignalChem; Tyro3 (aa 455-end): T22-11G, Axl (aa 473-end): A34-11G, Mer (aa 528-end): M51-112G) added to the (+) samples. The (-) sample had buffer only added in place of kinase. All samples were incubated at 37°C for 2 hours, and the reaction quenched by adding 0.5% TFA by reaction volume. The samples were desalted as previously described (above) and vacuum dried. Phosphopeptide enrichment was carried out according to manufacturer’s instructions. Samples were enriched first using High-Select™ TiO2 Phosphopeptide Enrichment Kit (ThermoFisher), and the flowthrough and wash fractions retained, dried, and enriched using the High-Select™ Fe-NTA Phosphopeptide Enrichment Kit (ThermoFisher).

### LC-MS/MS data acquisition

Samples were reconstituted in 0.1% formic acid (FA) in H_2_O and loaded onto a Dionex UltiMate 3000 RSlCnano system and run over a linear gradient (5-30% acetonitrile; 65 minutes) with a flow rate of 300 nL/min into LTQ Orbitrap Velos Mass Spectrometer (ThermoFisher). The MS was operated using data dependent mode with a resolution of 30,000 ppm with a scan range of 380 – 1800 m/z. MS/MS triggered for the top six abundant ions using high collision dissociation (HCD) and mass analyzer parameters set between 2 and 7 charge states.

### Data processing

Data was processed as previously described.^25^ In brief, peptides were identified using PEAKS Studio Xpro (Bioinformatics Solutions Inc.) and exported protein-peptides.csv lists were used for input into the PEAKS ModExtractor^35^, which combines all the input lists into a concatenated output file, containing a modification-centered list of all peptides. Tables S1-3 show a summary of the total number of unique peptides and phosphorylated peptides with A-score ≥ 30 identified for each kinase and replicate.

These data were input into KINATEST-ID V2.1.0 (https://github.com/llparkerumn/KINATESTIDv2) and filtered to include only nonredundant substrates found in all three kinase reaction replicates for each kinase, but not in any of the controls (Tyro3: n = 425; Axl: n = 270; Mer: n = 252). These filtered substrates were used as input to generate the Fisher Odds position-specific scoring matrix (PSSM) for each kinase. The candidate amino acids at each position were then permuted to generate the in-silico library of putative substrates. Each peptide sequence was given a score, calculated based on the odds ratio of each amino acid in the sequence. For each kinase, receiver-operator curve (ROC) analysis was performed using the input peptides lists to establish a threshold value for predicted peptide activity. Sequences with scores above this value are predicted to be substrates for the target kinases. Final synthetic substrates were scored and normalized as previously described.^25^ All KINATEST-ID V2.1.0 outputs are provided in the Supporting Information.

### Peptide synthesis and purification

Peptides were synthesized as previously described.^25^ In brief, the Symphony X Peptide Synthesizer (Protein Technologies) was used to synthesize peptides with an activation solution of HCTU in N-methylmorpholine (NMM) and dimethylformamide (DMF). 20% piperidine in DMF was used to deprotect Fmoc groups, and a cleavage cocktail of 94% trifluoroacetic acid (TFA), 2.5% H_2_O, 2.5% ethane dithiol and 1% Triisopropylsilane (TIS) used to cleave peptides from resin. Reverse phase HPLC (Agilent 1200 Series Infinity LCMS) was used to purify peptides to >90% purity. LC/MS characterization for all peptides is provided in the supporting information (Fig. S4).

### In vitro synthetic substrate assays with ELISA readout

Recombinant Tyro3, Axl, or Mer (SignalChem; Tyro3 (aa 455-end): T22-11G, Axl (aa 473-end): A34-11G, Mer (aa 528-end): M51-112G) was incubated with reaction mixture (25 mM HEPES pH 7.5, 10 mM MgCl2, 100 µM ATP, 1 µM Na_3_VO_4_ and 0.05 µg/µL BSA) to a final concentration of 10 nM for 15 minutes at room temperature, and the reaction started by adding peptide substrate to a final concentration of 20 µM. Timepoints were taken by quenching sample aliquots 1:1 in EDTA. The zero-minute timepoints were taken by adding substrate to pre-quenched kinase mixture. Quenched sample aliquots were incubated for 1 hour in streptavidin coated plates (ThermoFisher Scientific) in Tris-buffered saline with 0.05% Tween-20 (TBS-T) with 5% w/v milk. Following sample incubation, the wells were washed with TBS-T and incubated with 4G10 Anti-Phosphotyrosine Antibody horseradish peroxidase conjugate (Millipore Sigma; 1:5000 dilution in TBS-T 5% milk) for 1 hour. A solution of 0.1 mM Amplex Red (ThermoFisher Scientific) and 2.225 mM hydrogen peroxide in 50 mM sodium phosphate buffer, pH 7.4 was added to the wells and incubated in the dark for 30 minutes. Fluorescent measurements were taken on a Synergy Neo2 plate reader (Biotek) with an excitation wavelength of 532 nm and emission wavelength of 590 nm.

### Synthetic substrate *in vitro* assay with HPLC-MS readout

Reactions (300 µL total volume) using recombinant kinase (Tyro3, Axl, or Mer) were performed in triplicate at 25°C in kinase reaction buffer (25 mM HEPES pH 7.5, 10 mM MgCl2, 100 µM ATP, 1 µM Na_3_VO_4_ and 0.05 µg/µL BSA) with a kinase concentration of 10 nM. After incubating for 15 minutes, reactions were initiated with the addition of pooled substrate (final concentration 20 µM). Reaction aliquots (50 µL) were withdrawn and quenched with 10 µL 20% TFA (3.3% TFA in the quenched solution). Sample aliquots (50 µL) were run on HPLC (Agilent 1200 series) using a C18 Agilent Zorbax column (2.1 x 250mm, 5-micron) at 0.25 mL/min with acetonitrile increasing from 5% to 20% over 90 min. Reaction progress was analyzed via MS (Agilent 6130A) through integration of extracted ion chromatograms (EIC) corresponding to products (P) and substrates (S) (Agilent ChemStation) and normalized according to equation:

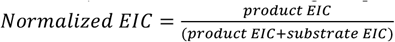

Multiple products were observed for Axl substrates Syn A, Syn C and Syn D. The phosphorylation sites of the products at different retention times were determined using MS/MS as shown in the supporting information (Fig. S5).

### AlphaFold2 modeling

In our study, we utilized the colabfold_batch command line application from the ColabFold package (version 1.5.1, commit b4c1bc7) for structural modeling. Initially, we constructed 3D structures of three kinases without peptides. The kinase domain sequences were extracted from UniProt (Tyro3_Q06418 - 518-788, Axl_P30530 - 536-805, and MER_Q12866 - 587-856) and provided to AlphaFold2 for predictions. To generate a diverse set of conformations for the kinase structures, we followed the MSA subsampling protocol while maintaining the same ratio of ‘max_msa_clusters’ and ‘max_extra_msa’ parameters. We used the alphafold2_ptm models (all five neural networks were included in calculations) with 24 recycles, an early stop tolerance of zero, and one random seed equal to zero. We saved all intermediate structures generated during recycling and selected the most confident model for each kinase based on pLDDT and pTM scores across all structures with catalytically active conformations. We used KinCore (https://github.com/vivekmodi/Kincore-standalone) to define the kinase conformations, considering them catalytically active if the program defined spatial label equal DFGin, αC-helix label equal Chelixin, and dihedral label as BLAminus.

Next, we employed the selected kinase models as templates for complex prediction of kinase-peptide pairs. We concatenated the catalytic domain sequence of the kinase with the peptide sequence with a colon symbol between them. We used the alphafold2_multimer_v3 model with 24 recycles, an early stop tolerance of zero, and one random seed equal to 42. The MSA mode was presented in single sequence and unpaired mode. We saved only structures predicted on the last iteration of recycling by the five neural networks implemented in the alphafold2_multimer_v2 model.

We calculated the predicted class for the resulting models based on two parameters. The first parameter was the minimal distance between the C-*γ* atom of Asp in HRD motif in the catalytic loop of the kinase and the hydroxyl oxygen of any Tyr in the peptide sequence. If the distance was less than 5 Å, we considered that the peptide had the correct position relative to the kinase domain. The second parameter was the class of kinase conformation from KinCore. During predictions AF2 produces 5 different structural models for each kinase-peptide pairs, and we treated a given kinase-peptide pairing as a true substrate if any of the 5 models met the required criteria.

## Supporting information

Structure models

Phosphoproteomics summary data

KINATESTID files

GraphPad Prism and Excel files

## SUPPORTING INFORMATION

Raw data, PEAKS X Studio search results files, and PEAKS-ModExtractor output files are available at the MassIVE repository (<u>doi:10.25345/C5513V612</u>).

“Supporting information.pdf” includes additional data figures and peptide characterization.

“Tables S1-S3. KALIP unique peptide summaries.zip” (zipped folder) contains Excel files summarizing the phosphopeptides observed from the KALIP reactions for each kinase.

“KINATESTIDV2.1.0.zip” (zipped folder) contains the output files for each kinase’s KALIP dataset from KINATESTIDV2.1.0 analysis in R.

“GraphPad Prism and Excel analyses.zip” (zipped folder) contains files from data analyses that were used in figures in the main manuscript and the supporting information, for Figures 2-4, Fig. S1 and Table S4.

“ActiveModels.zip” contains structural data files for each AF2 output model of productive substrate-enzyme interaction.

## Author Contributions

NEW and LLP conceived the project, with contributions on structural modeling from GA and JK. NEW, CH, and JLH performed biochemical and LC/MS experiments. GA performed structural modeling analyses. NEW, CH, JLH, and GA analyzed data. NEW, GA, JK, and LLP wrote the manuscript.

## ACKNOWLEDGMENT

We appreciate mass spectrometry analysis assistance and data analysis support from Peter Villalta and Yingchun Zhao (Masonic Cancer Center MS Core Facility), and LeeAnn Higgins (Center for Mass Spectrometry and Proteomics). We also appreciate copy editing assistance from Milton Andrews. Funding was provided by T32CA009138. (NEW), R01GM141513 and P30CA006927 (JK), and R01GM146386 and R01CA182543 (LLP).

## ABBREVIATIONS

KALIP: kinase assay linked with phosphoproteomics
LC-MS/MS: liquid chromatography tandem mass spectrometry
HPLC: high performance liquid chromatography.

## TOC Graphic

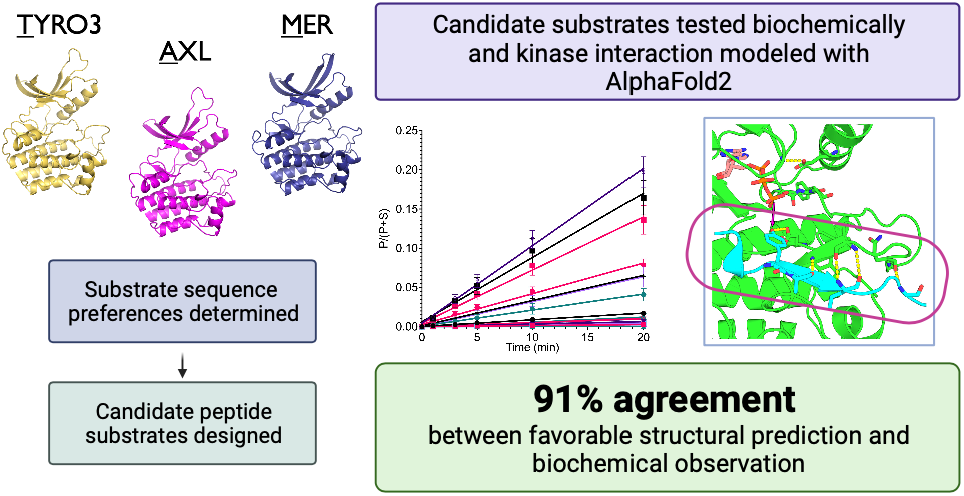

## Notes

https://massive.ucsd.edu/ProteoSAFe/dataset.jsp?accession=MSV000092784

### Competing Interest Statement

The authors have declared no competing interest.

