## Supplementary figures and images for "Novel substrate prediction for the TAM family of RTKs using phosphoproteomics and structure-based modeling"

### AXL_heatmap.png

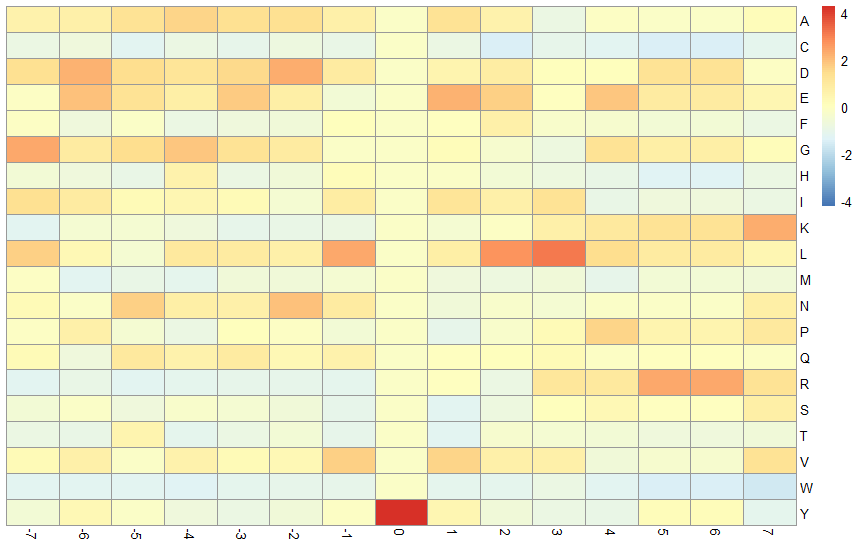

### AXL_venn.png

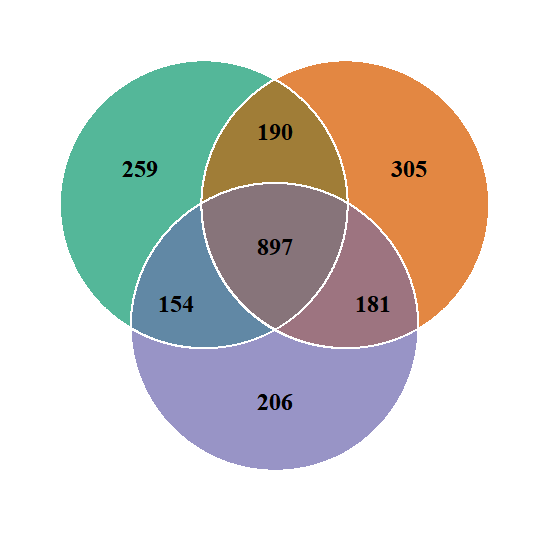

### MER_528-999_heatmap.png

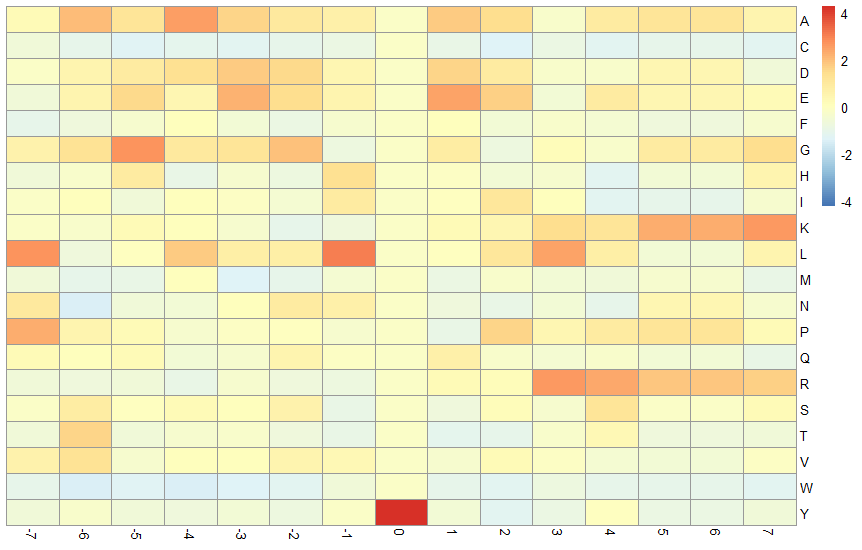

### MER_528-999_venn.png

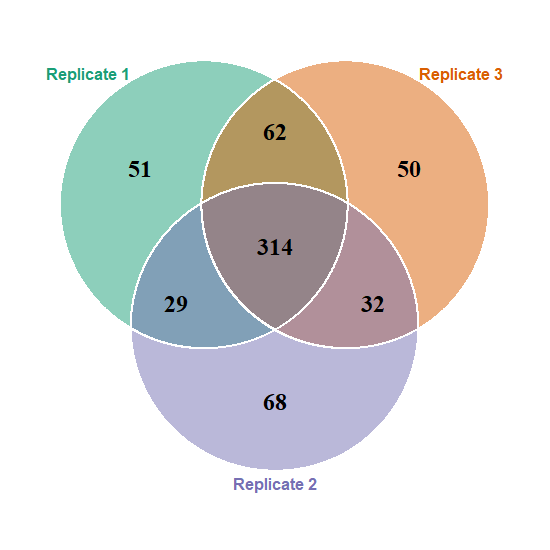

### roc.jpg

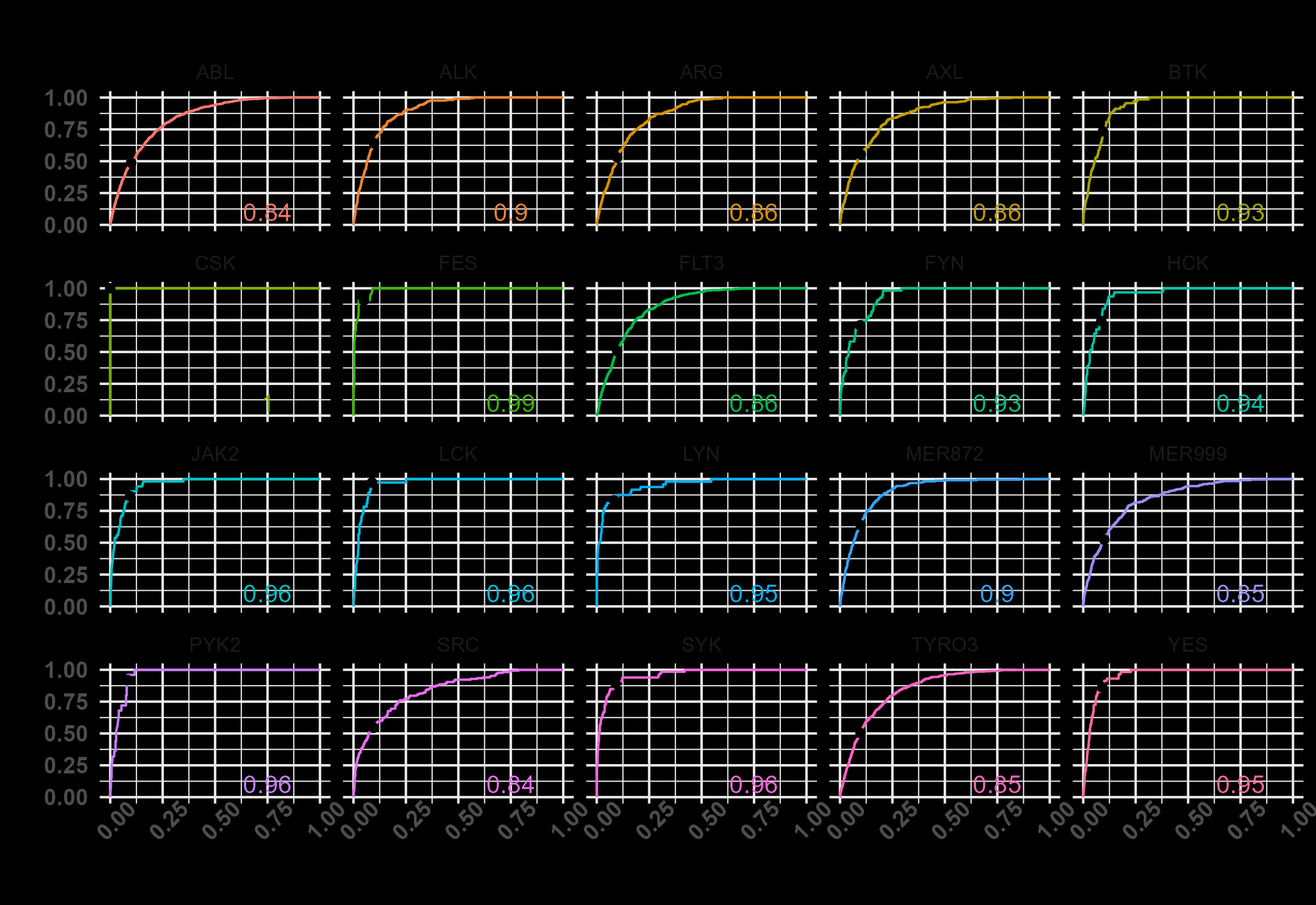

### roc.jpg

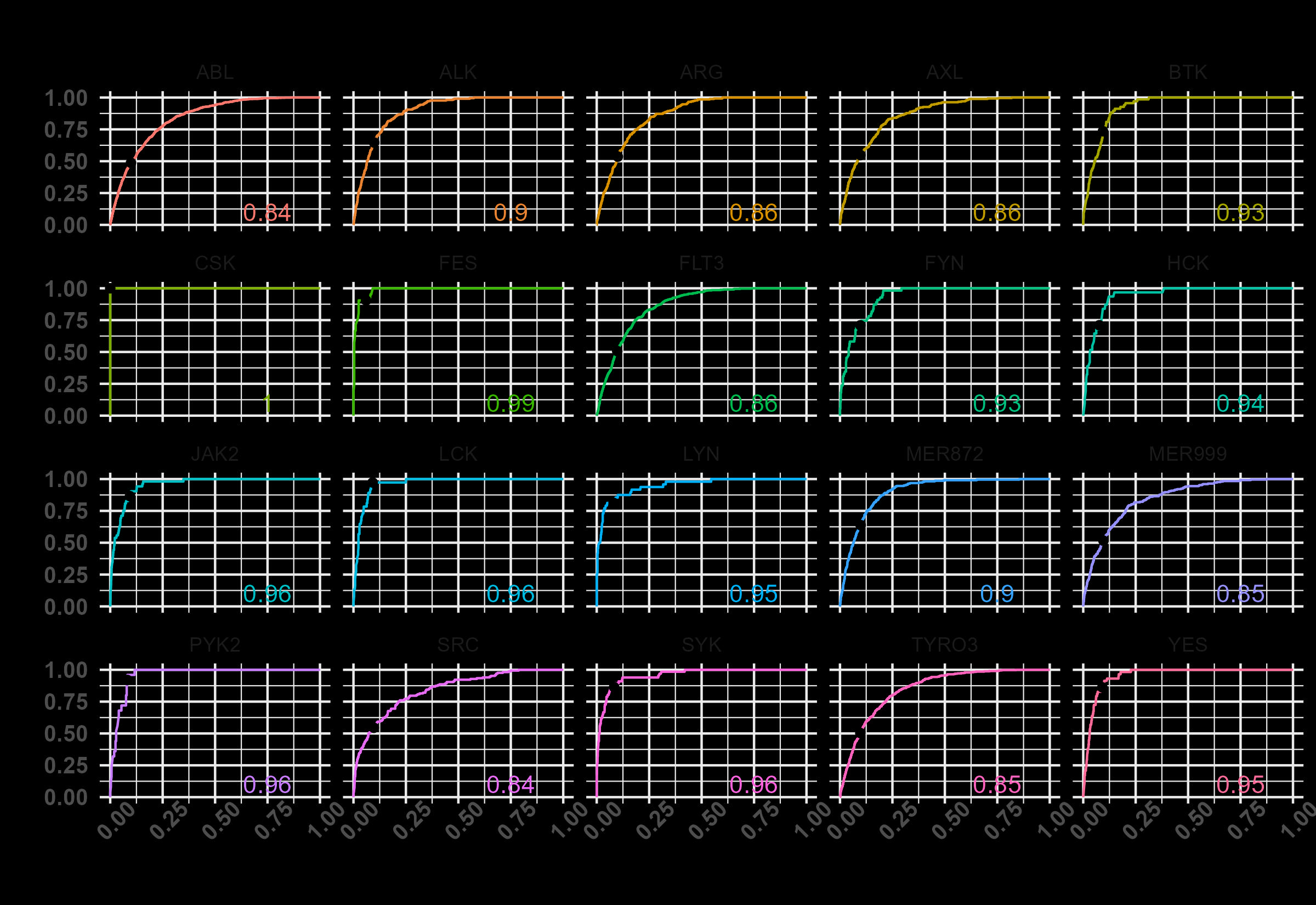

### roc.jpg

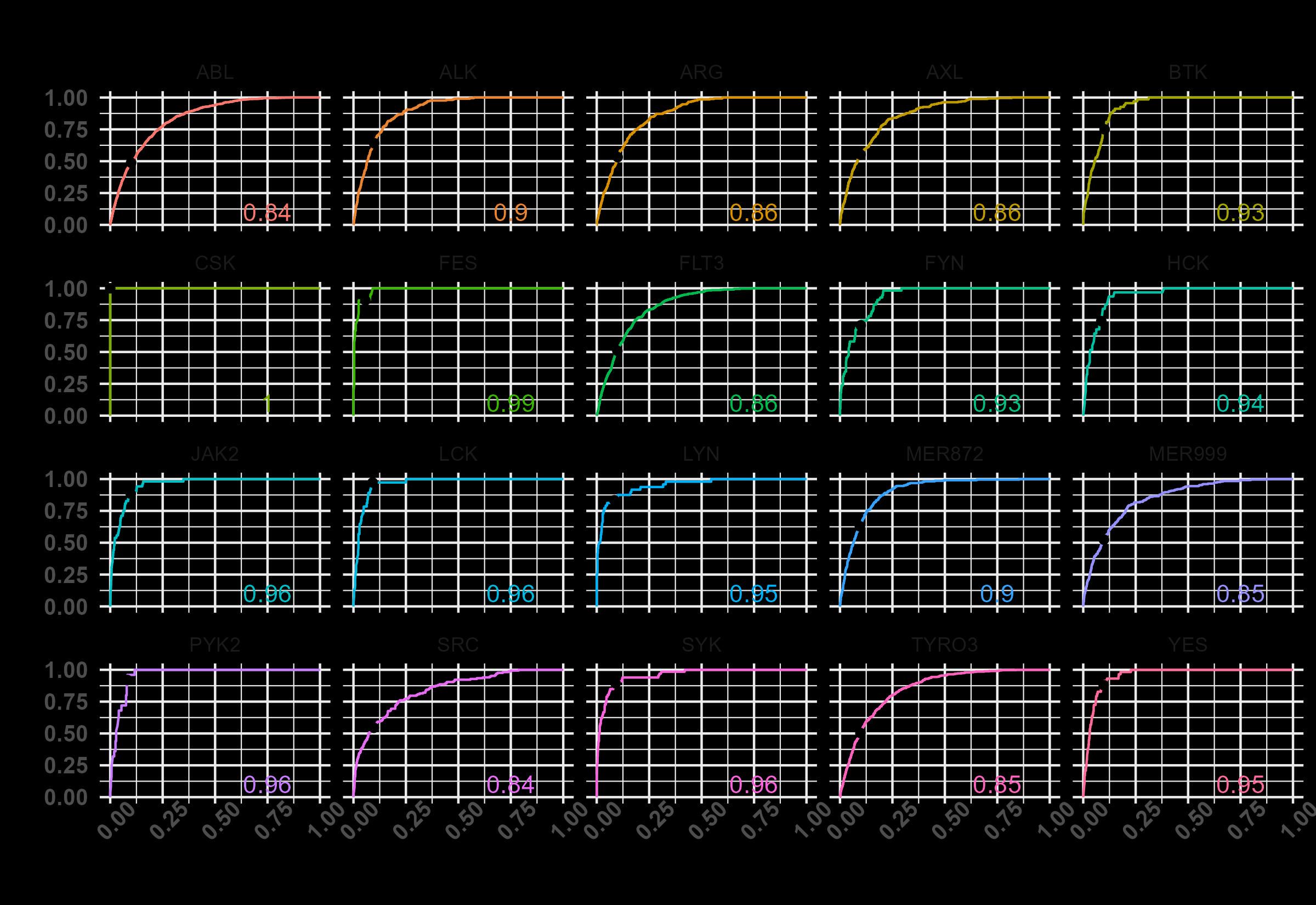

### screener_cutpoints.jpg

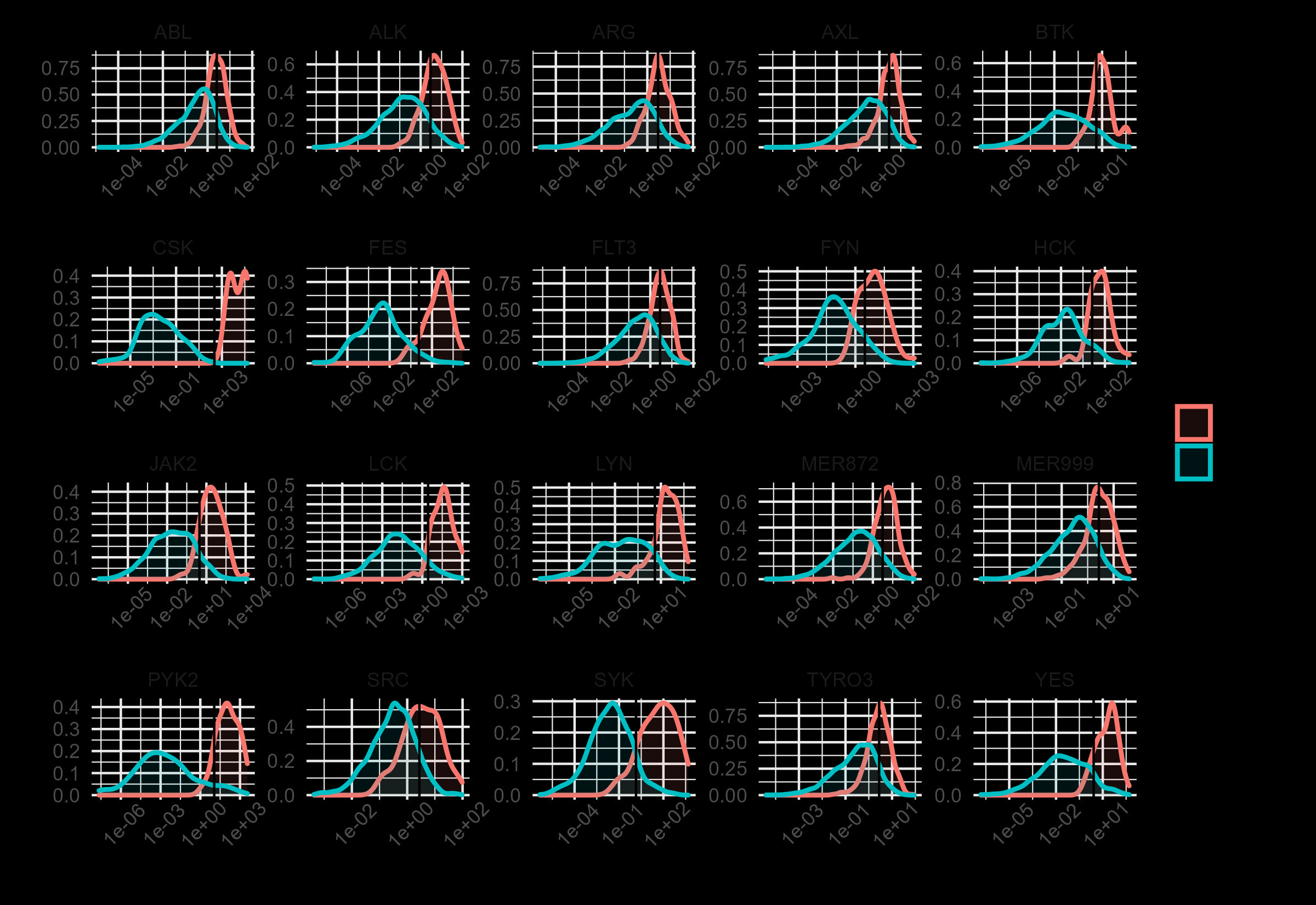

### screener_cutpoints.jpg

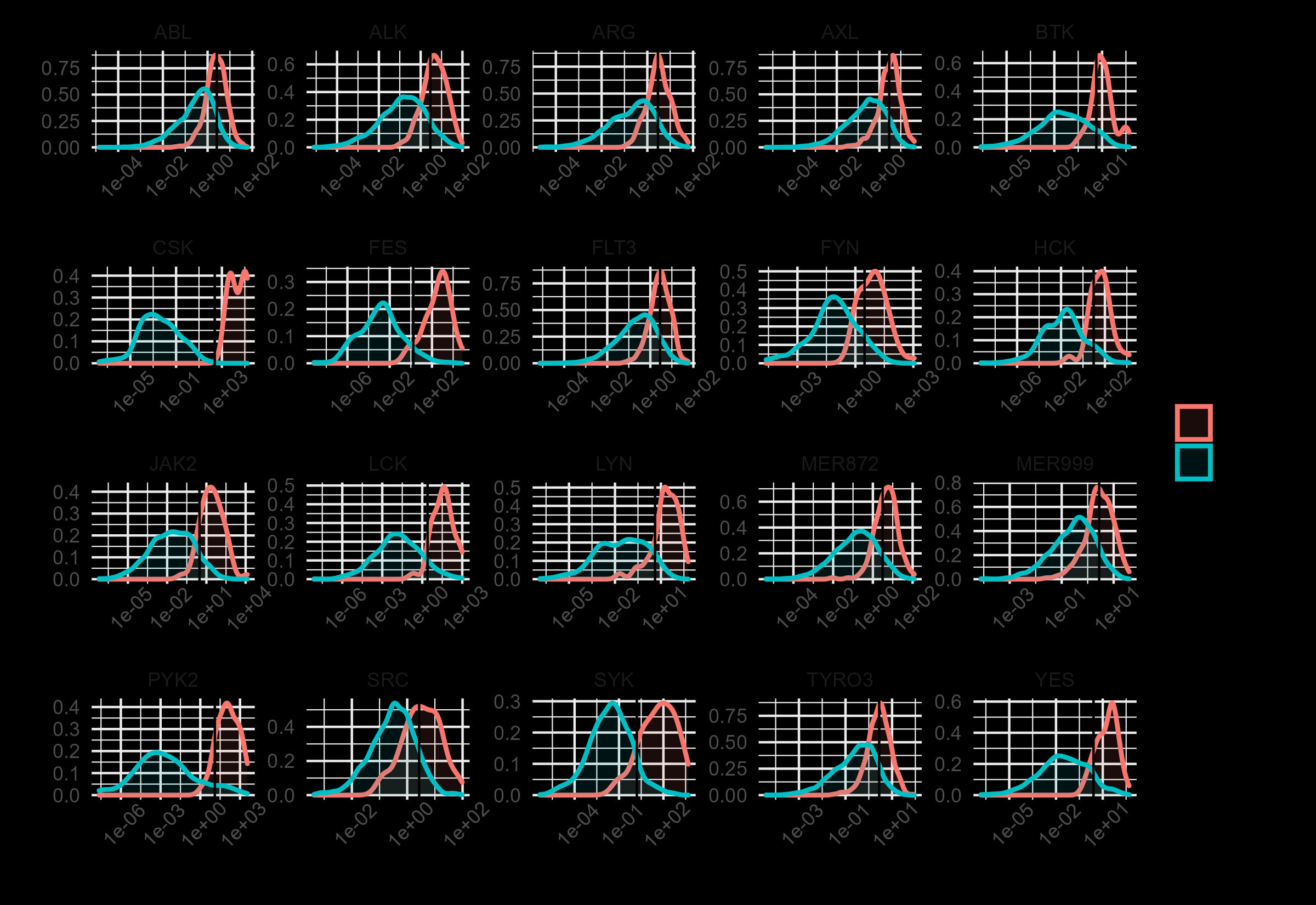

### screener_cutpoints.jpg

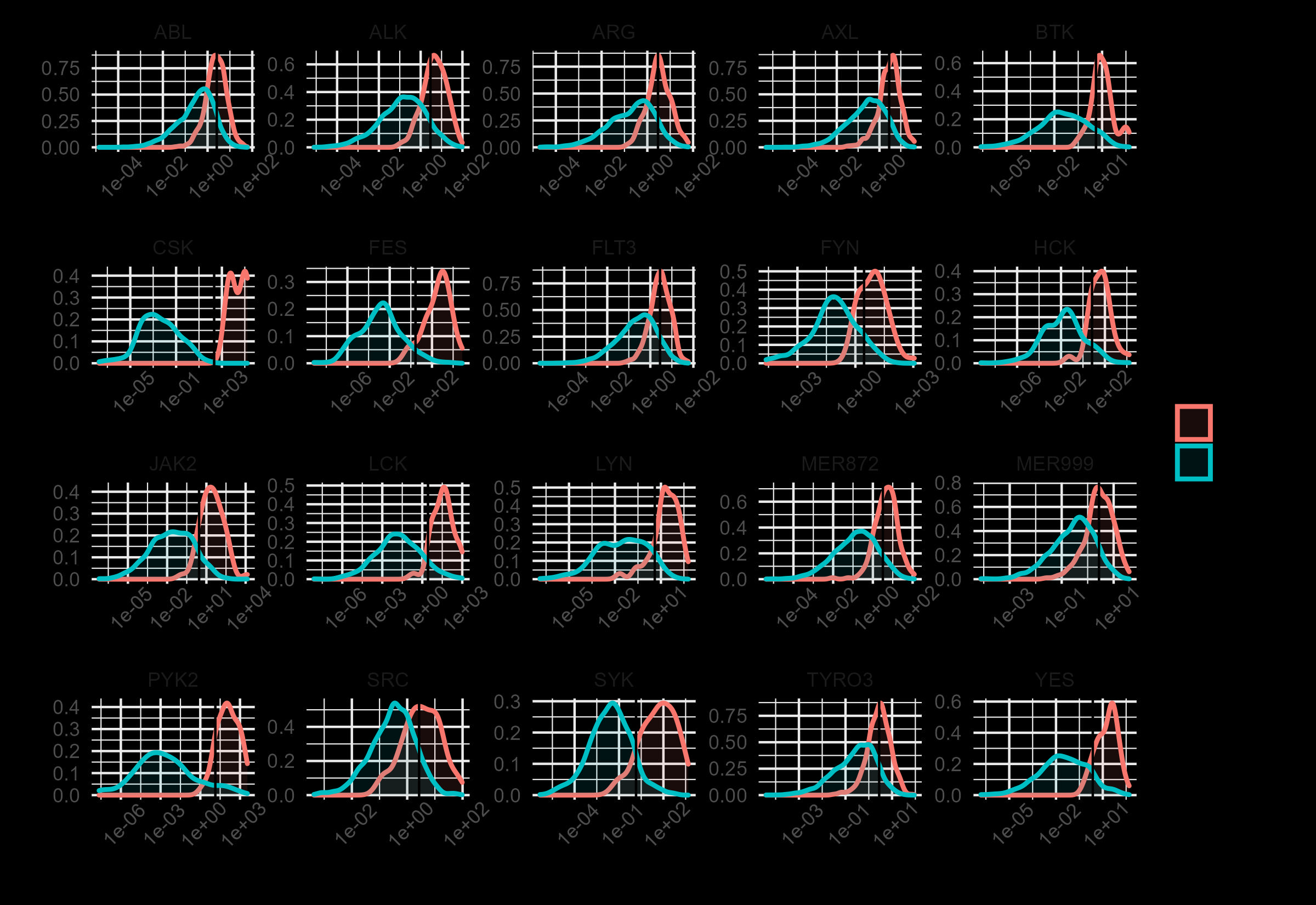

### TYRO3_heatmap.png

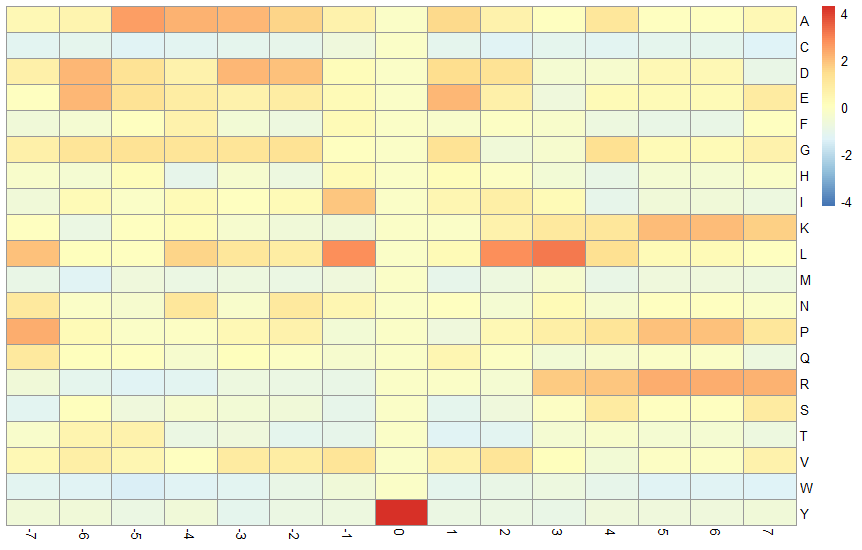

### TYRO3_venn.png

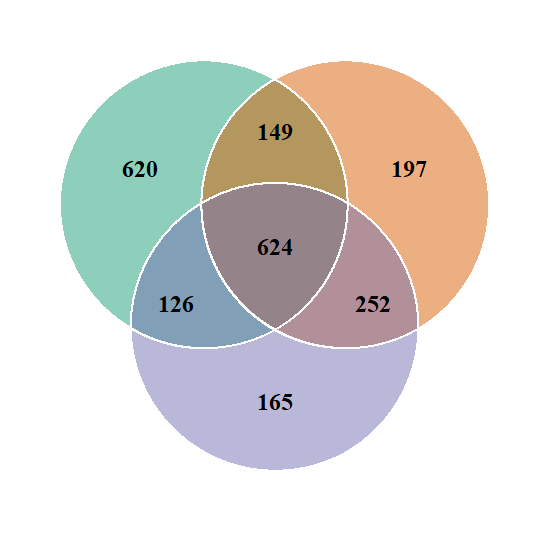
