## Supporting Info document for "Novel substrate prediction for the TAM family of RTKs using phosphoproteomics and structure-based modeling"

Supporting Information for

### Prediction of novel synthetic peptide substrates for the TAM family of RTKS using phosphoproteomics and structural modeling

Naomi E. Widstrom,[1] Grigorii V. Andrianov,[2] Jason L. Heier,[1] Celina Heier,[1] John Karanicolas,[2] Laurie L. Parker[1]

[1] Department of Biochemistry, Molecular Biology and Biophysics, College of Biological Sciences, University of Minnesota Twin Cities, Minneapolis, MN 55455

2 Cancer Signaling & Microenvironment Program, Fox Chase Cancer Center, Philadelphia, Pennsylvania 19111-2497

#### Table of Contents:

*Figure S1. Characteristics of N- and C-terminal length of phosphopeptides from in vitro phosphorylation reactions with Tyro3, Axl, and Mer respectively.* 2

*Figure S2. Progress curves for phosphorylation of Universal Substrate 5 (U5: DEAIYATVAGGK$_{biotin}$GG) by the TAM family kinases, confirming kinase activity for enzymes used in this work.* 3

*Figure S3. Plots of product phosphopeptide (P) and substrate (S) EIC over time in pooled substrate experiments for Tyro3, Axl and Mer.* 4

*Figure S4. Substrate peptide characterization by LC/MS* 5

Tyro3 synthetic substrate A (EGLYHHRNHPGGK$_{Biotin}$GG). 5
Tyro3 synthetic substrate B (HTIYHHKNHPGGK$_{Biotin}$GG) 5
Tyro3 synthetic substrate C (HQNYDHKNHPGGK$_{Biotin}$GG) 6
Tyro3 synthetic substrate D (HQNYTHKNPRGGK$_{Biotin}$GG) 6
Tyro3 synthetic substrate E (HGHYGHPNHPGGK$_{Biotin}$GG) 7
Tyro3 synthetic substrate F (HQNYTHKNPPGGK$_{Biotin}$GG) 7
Axl synthetic substrate A (NDENNYYYRGGRGGK$_{Biotin}$GG) 8
Axl synthetic substrate B (NDENNYAFRGGRGGK$_{Biotin}$GG) 8
Axl synthetic substrate C (NDENYYYYRGGRGGK$_{Biotin}$GG) 9
Axl synthetic substrate D (NDENYYYYTGGRGGK$_{Biotin}$GG) 9
Mer synthetic substrate A (NEGKHGHYAILKDDRGGK$_{Biotin}$GG) 10
Mer synthetic substrate B (NEGKHGFYDARKDDKGGK$_{Biotin}$GG) 10
Mer synthetic substrate C (NFAKHGFYDARRADRGGK$_{Biotin}$GG) 11
Mer synthetic substrate D (NFGEEGFYAARKEDKGGK$_{Biotin}$GG) 11
Mer synthetic substrate E (DHGHYAILPGGK$_{Biotin}$GG) 12
Universal peptide 5 (DEAIYATVAGGK$_{Biotin}$GG) 12

***Figure S5.*** *Identification of phosphorylation sites for Axl substrates containing multiple tyrosines and producing multiple products.* 13

*AXL Syn A:* NDENNYYYRGGRGGBGG 13
*AXL Syn C:* NDENYYYYRGGRGGBGG 15
*AXL Syn D:* NDENYYYYTGGRGGBGG 18

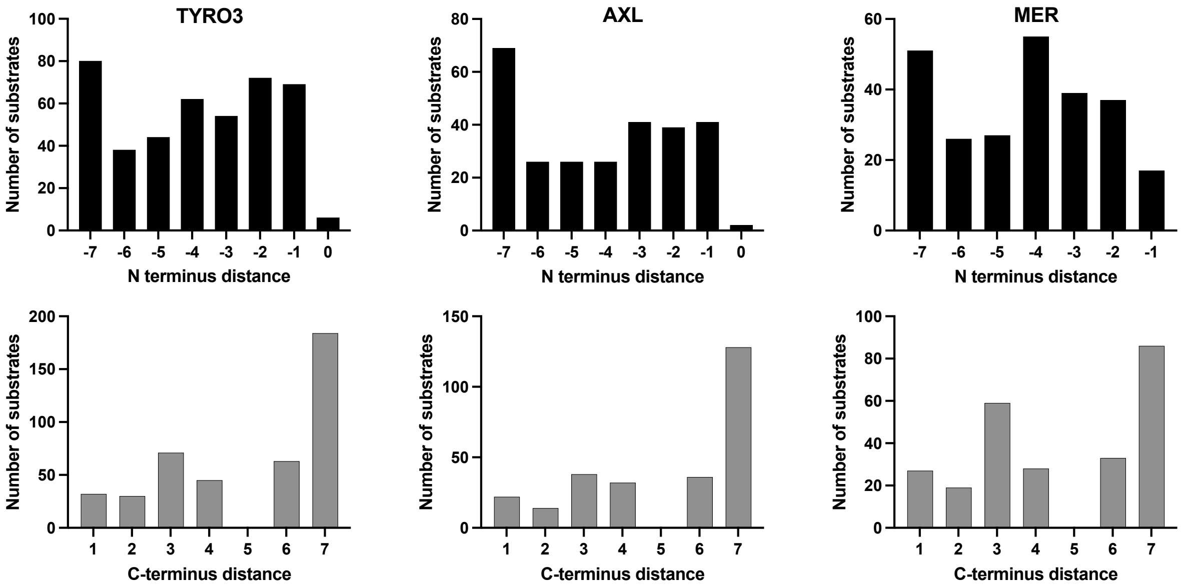


Figure S1. Characteristics of N- and C-terminal length of phosphopeptides from in vitro phosphorylation reactions with Tyro3, Axl, and Mer respectively.

The number of peptides observed with each given distance of the phosphosite from the terminus (-7 for N terminal distances, +7 for C-terminal distances) was counted and plotted as number of substrates per position, per terminus, for each kinase.

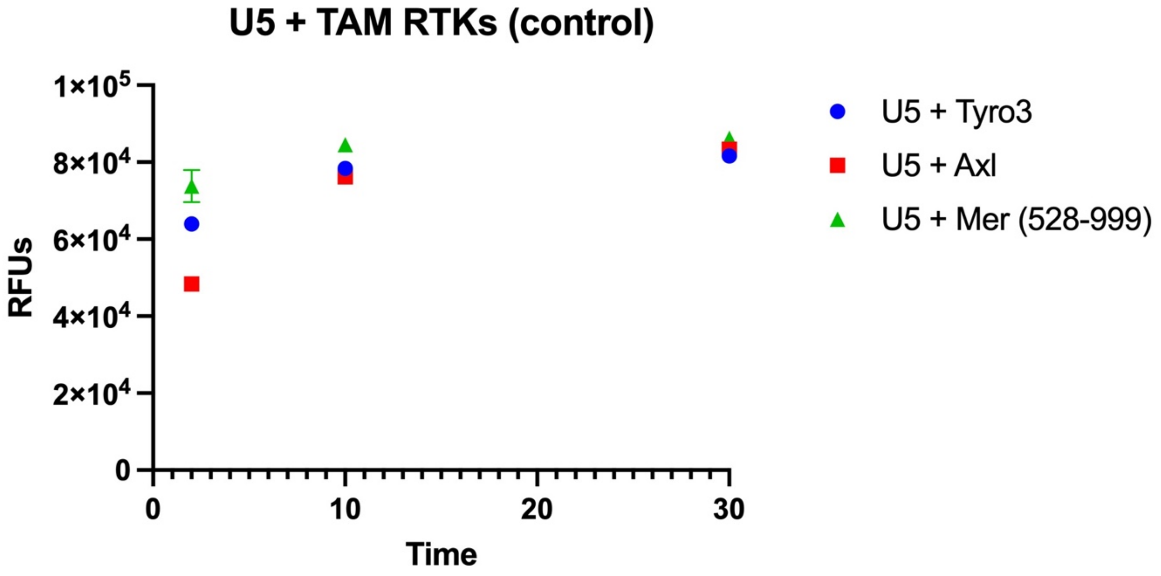


Figure S2. Progress curves for phosphorylation of Universal Substrate 5 (U5: DEAIYATVAGGK$_{biotin}$GG) by the TAM family kinases, confirming kinase activity for enzymes used in this work.

Assay conditions as described in the main manuscript Experimental section. Reaction aliquots quenched 1:1 in EDTA at timepoints 2, 10, 30 minutes to verify activity of the kinases used in the KALIP reactions. Data points average of 3 replicates ± SD. RFU, relative fluorescent units.

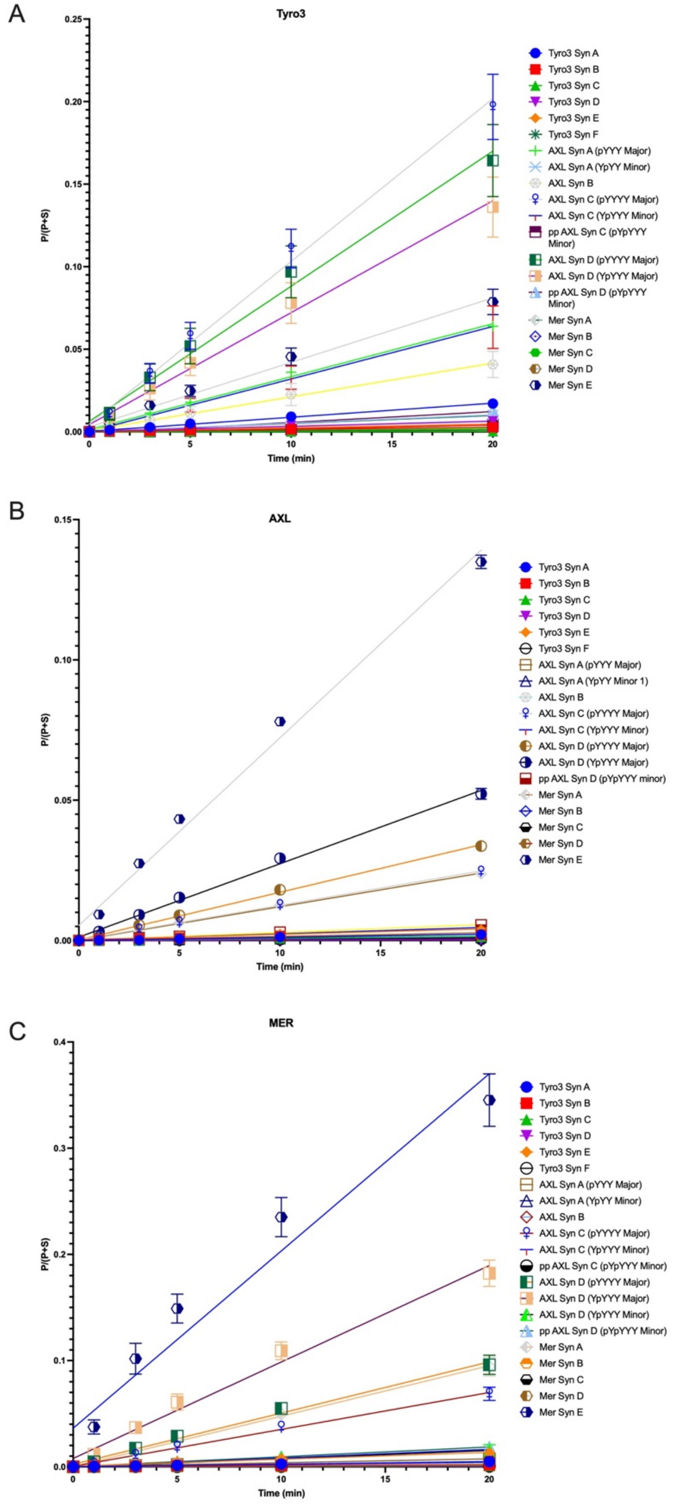


Figure S3. Plots of product phosphopeptide (P) and substrate (S) EIC over time in pooled substrate experiments for Tyro3, Axl and Mer.

See Supplemental File “Fig S3 LCMS TAM All substrate assay - product EIC vs time.pzfx” for full data and slope calculations.

Figure S4. Substrate peptide characterization by LC/MS.

Tyro3 synthetic substrate A (EGLYHHRNHPGGK$_{Biotin}$GG).

- $[M + H]^{1+}$ = 1841
- $[M + 2H]^{2+}$ = 921

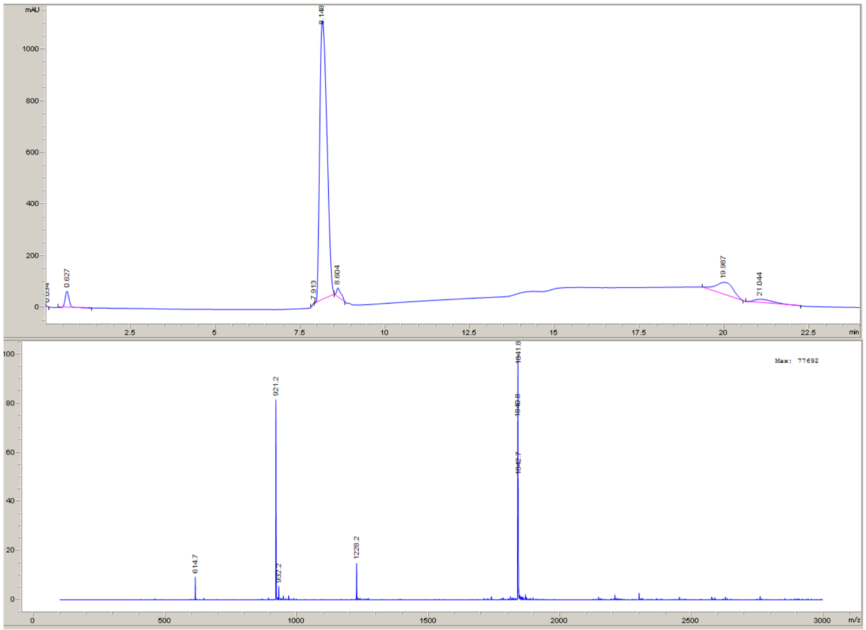


Tyro3 synthetic substrate B (HTIYHHKNHPGGK$_{Biotin}$GG)

- $[M + H]^{1+}$ = 1865
- $[M + 2H]^{2+}$ = 933

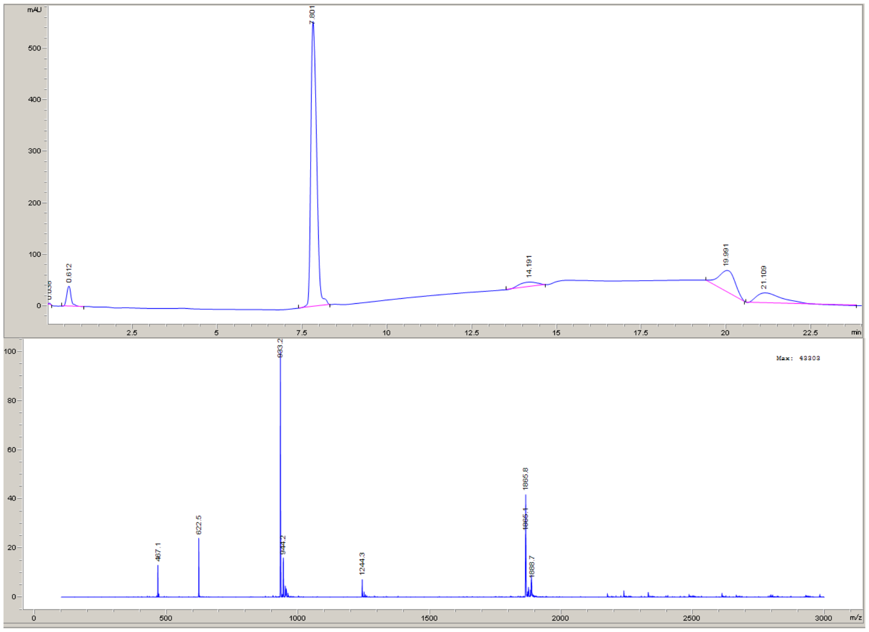

Tyro3 synthetic substrate C (HQNYDHKNHPGGK$_{Biotin}$GG)

- $[M + H]^{1+}$ = 1871
- $[M + 2H]^{2+}$ = 936

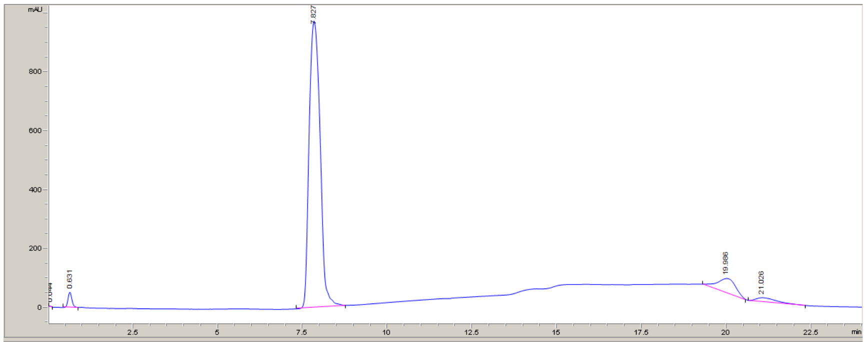


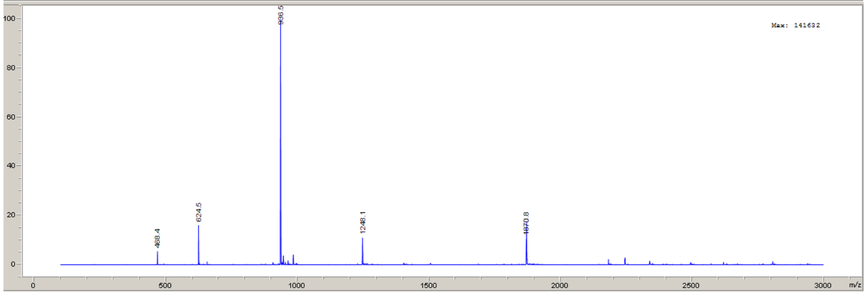


Tyro3 synthetic substrate D (HQNYTHKNPRGGK$_{Biotin}$GG)

- $[M + 1H]^{1+}$ = 1876
- $[M + 2H]^{2+}$ = 938

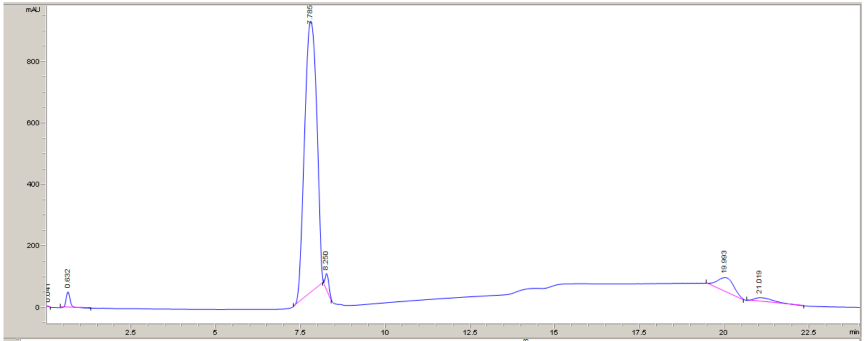


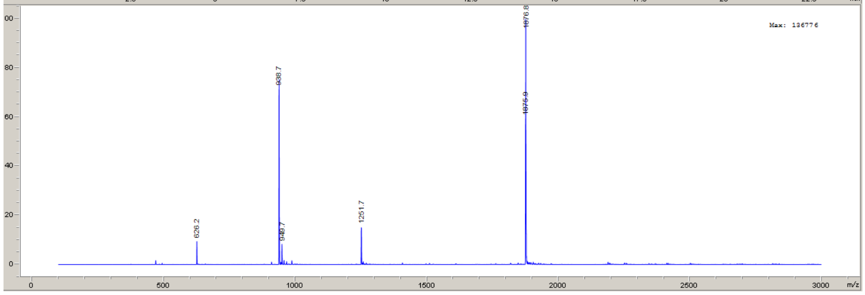

#### Tyro3 synthetic substrate E (HGHYGHPNHPGGK$_{Biotin}$GG)

- $[M + H]^{1+} = 1733$
- $[M + 2H]^{2+} = 867$

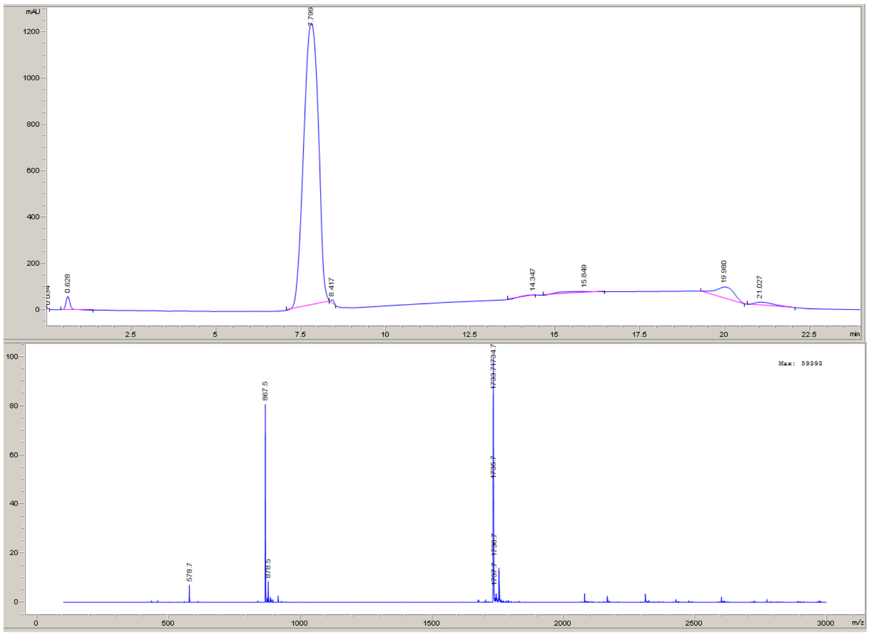


#### Tyro3 synthetic substrate F (HQNYTHKNPPGGK$_{Biotin}$GG)

- $[M + H]^{1+} = 1817$
- $[M + 2H]^{2+} = 909$

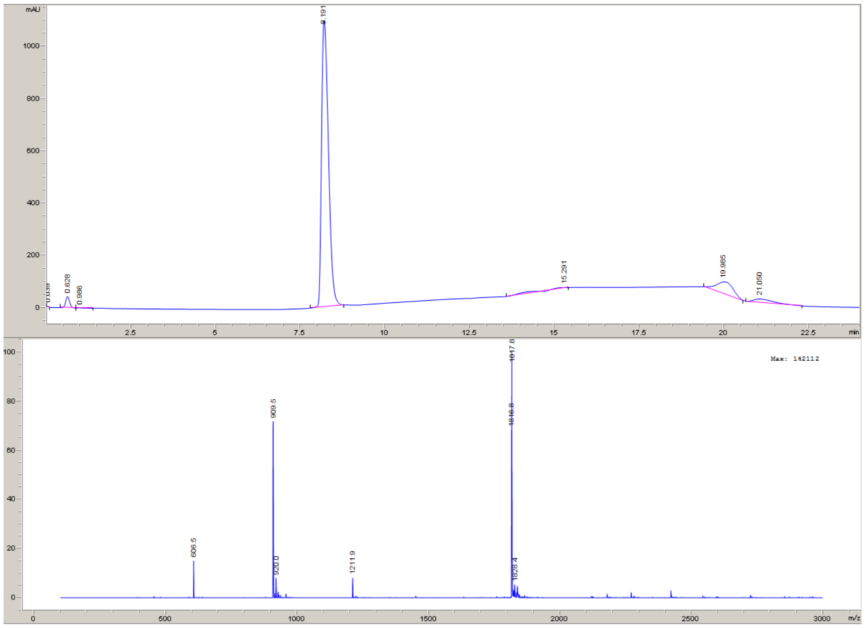

Axl synthetic substrate A (NDENNYYYRGGRGGK$_{Biotin}$GG)

- $(M + 2H)^{2+}$ = 1052
- $(M + 3H)^{3+}$ = 701

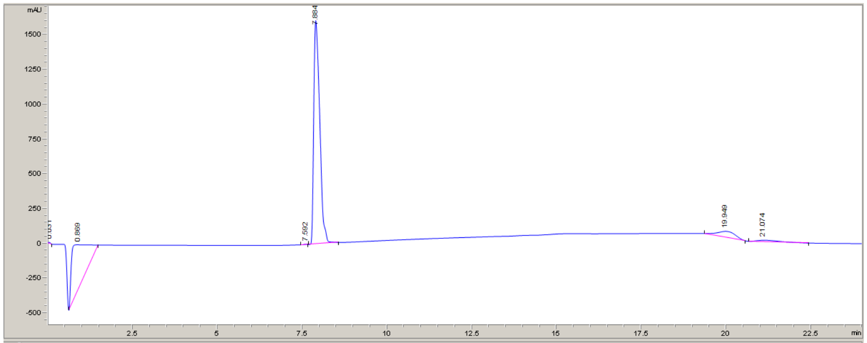


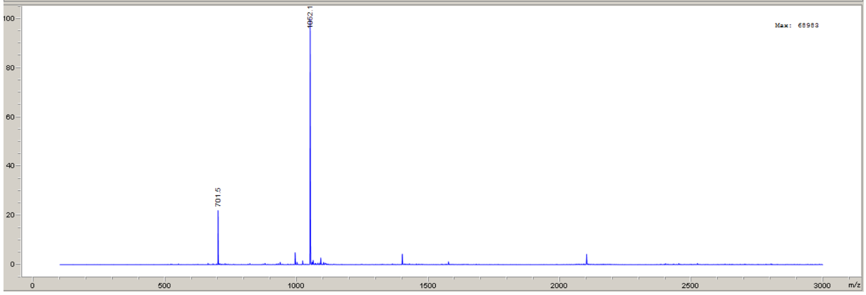


Axl synthetic substrate B (NDENNYAFRGGRGGK$_{Biotin}$GG)

- $[M + 2H]^{2+}$ = 998

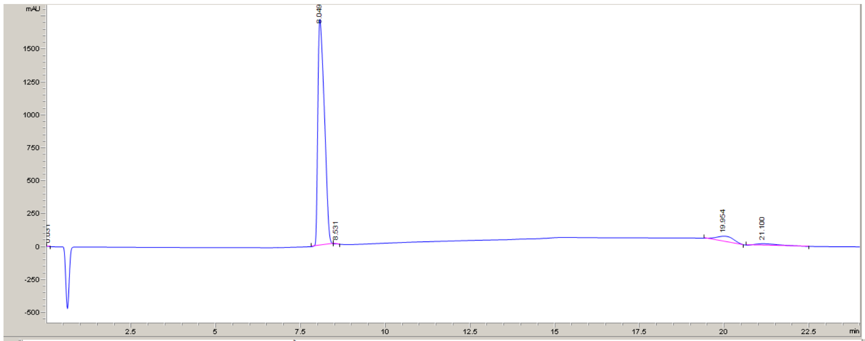


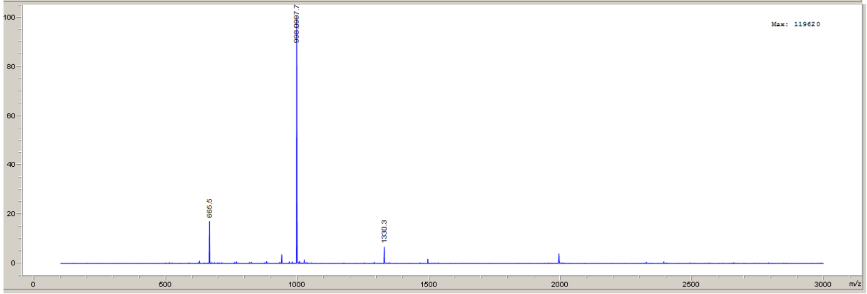

Axl synthetic substrate C (NDENYYYYRGGRGGK$_{Biotin}$GG)

- $[M + 2H]^{2+} = 1076$
- $[M + 3H]^{3+} = 718$

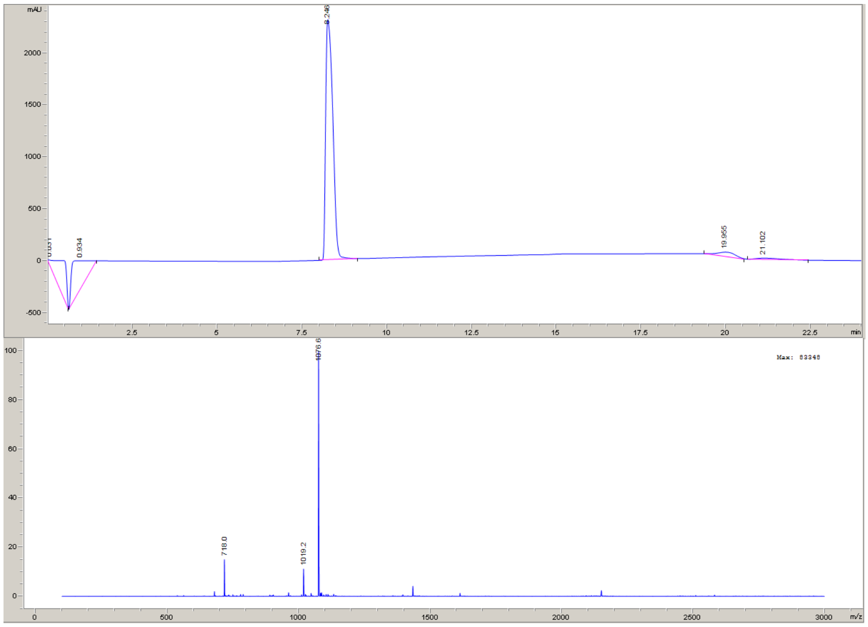


Axl synthetic substrate D (NDENYYYYTGGRGGK$_{Biotin}$GG)

- $(M + H)^{1+} = 2097$
- $(M + 2H)^{2+} = 1049$

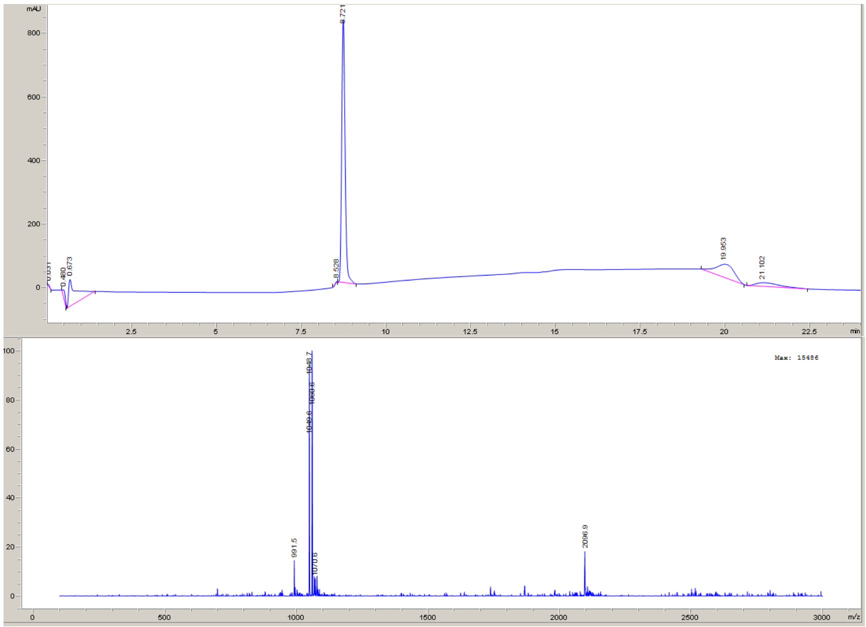

#### Mer synthetic substrate A (NEGKHGHYAILKDDRGGK$_{Biotin}$GG)

- $[M + 2H]^{2+}$ = 1167.8
- $[M + 3H]^{3+}$ = 779

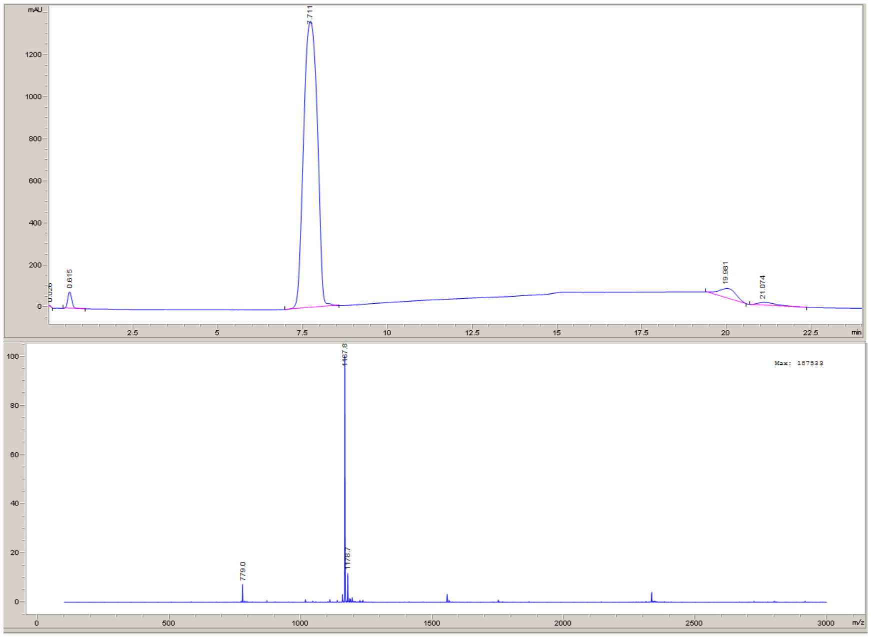


#### Mer synthetic substrate B (NEGKHGFYDARKDDKGGK$_{Biotin}$GG)

- $[M + 2H]^{2+}$ = 1181

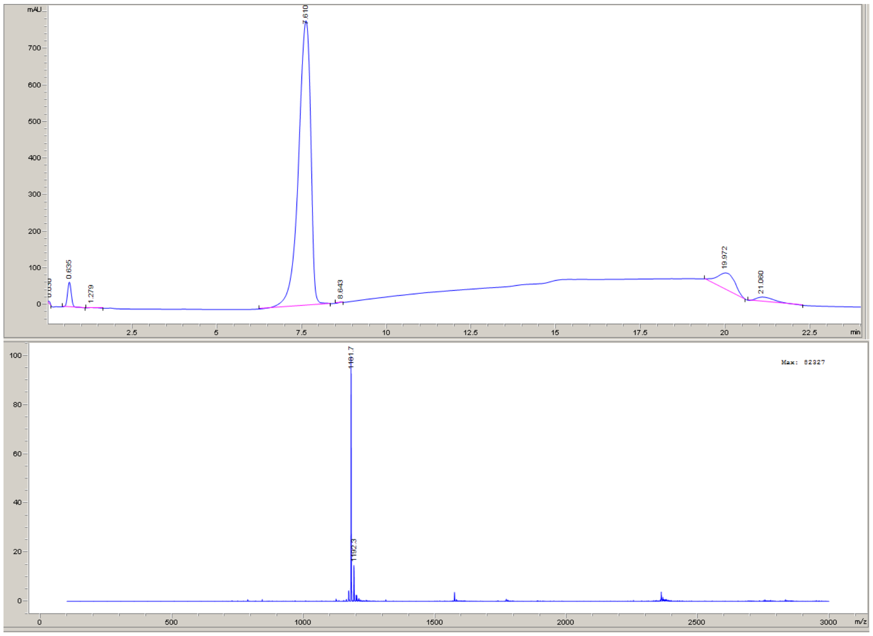

Mer synthetic substrate C (NFAKHGFYDARRADRGGK$_{Biotin}$GG)

- $[M + H]^{1+} = 2405.7$
- $[M + 2H]^{2+} = 1203.7$

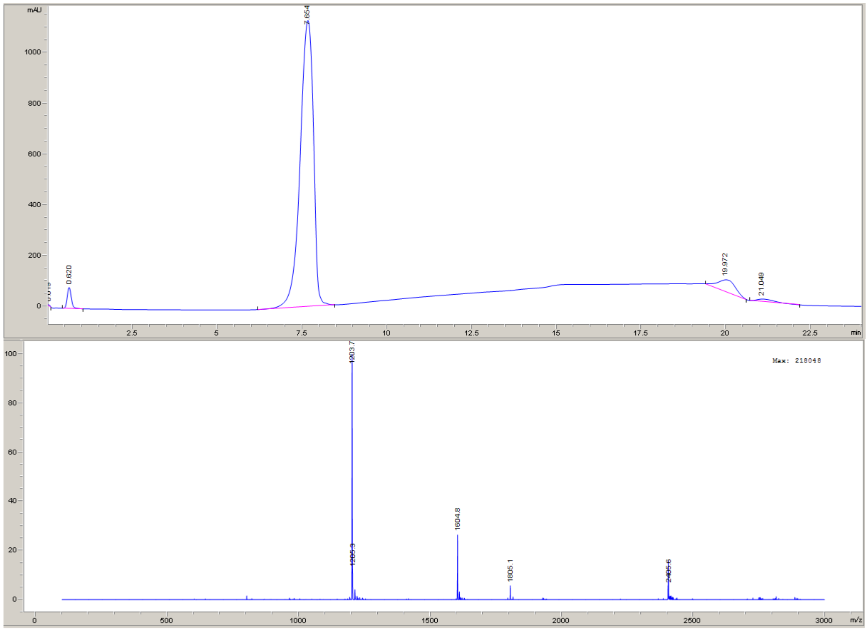


Mer synthetic substrate D (NFGEEGFYAARKEDKGGK$_{Biotin}$GG)

- $[M + 2H]^{2+} = 1172$

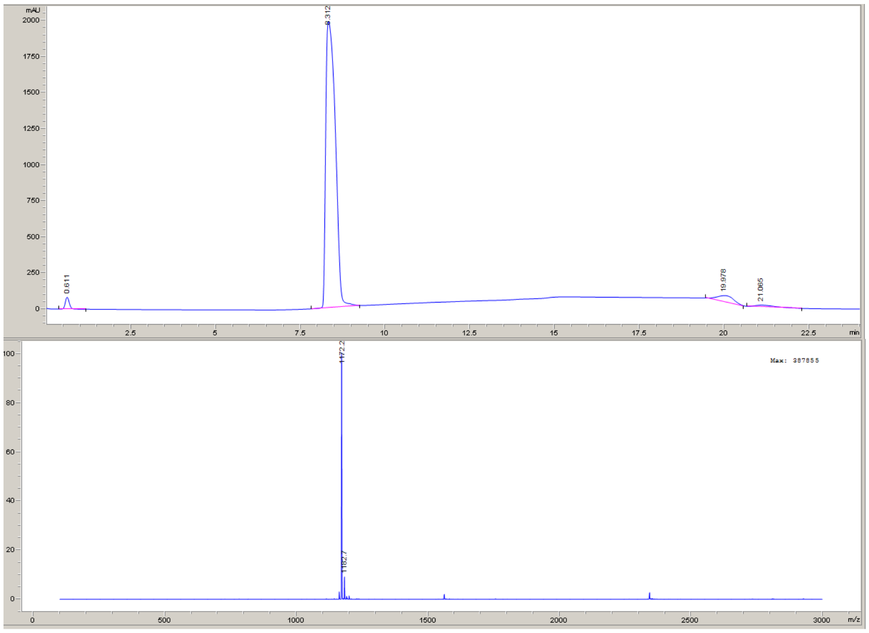

#### Mer synthetic substrate E (DHGHYAILPGGK$_{Biotin}$GG)

- $[M + H]^{1+}$ = 1603.8

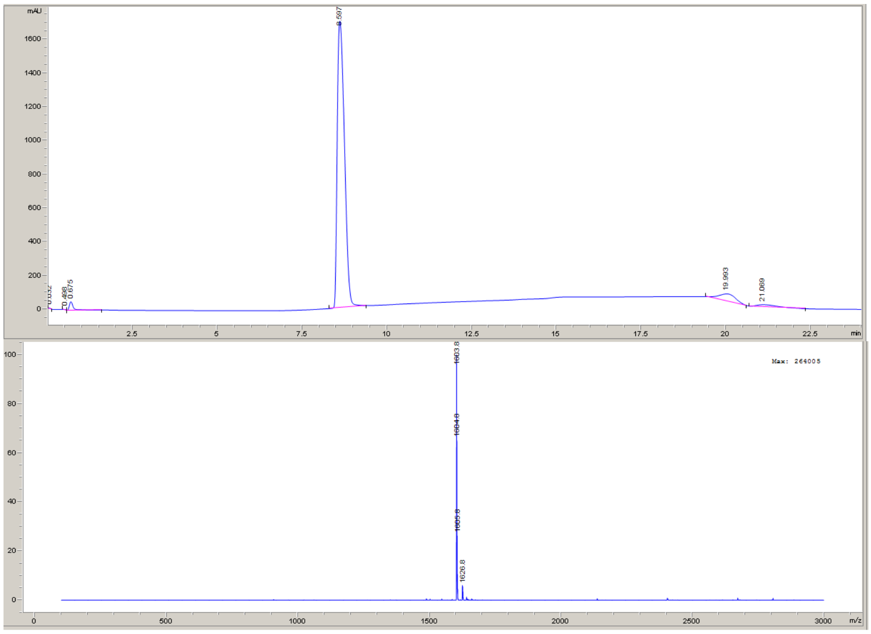


#### Universal peptide 5 (DEAIYATVAGGK$_{Biotin}$GG)

- $[M + H]^{1+}$ = 1534

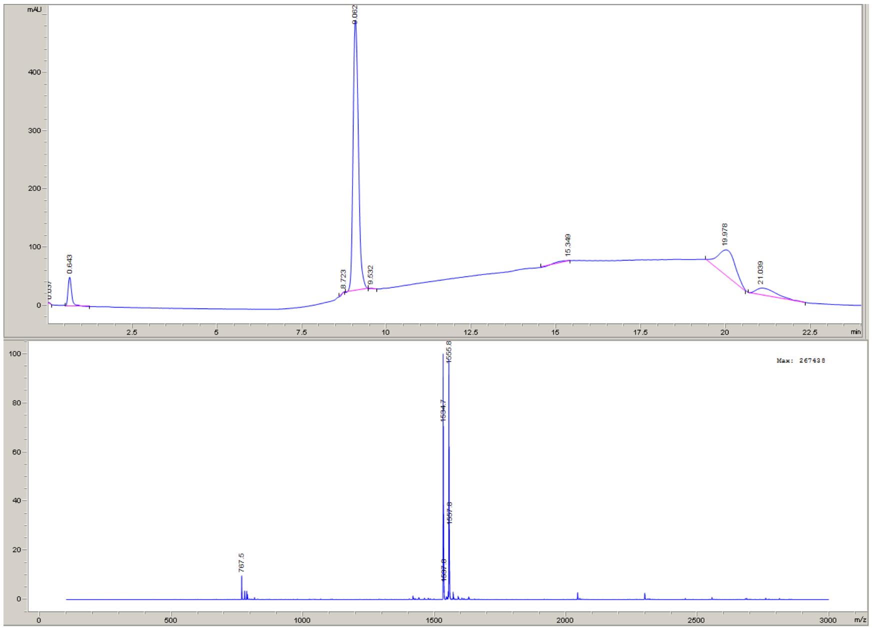

**Figure S5.** Identification of phosphorylation sites for Axl substrates containing multiple tyrosines and producing multiple products.

*AXL Syn A:* NDENNYYYRGGRGGBGG

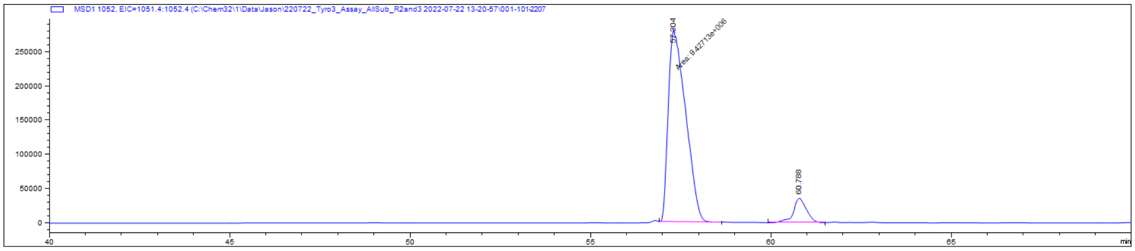

AXL Syn A at assay **t = 0 min; EIC of 1052**

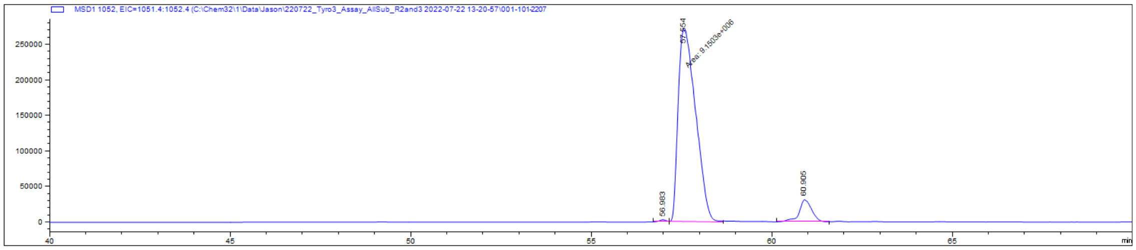

AXL Syn A at assay **t = 20 min; EIC of 1052**

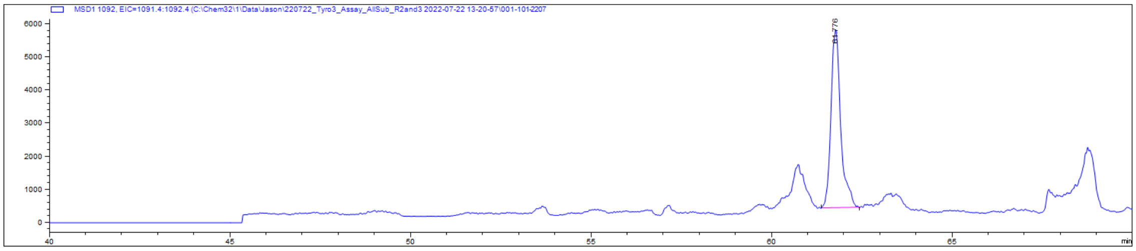

pAXL Syn A at assay **t = 0 min; EIC of 1092**

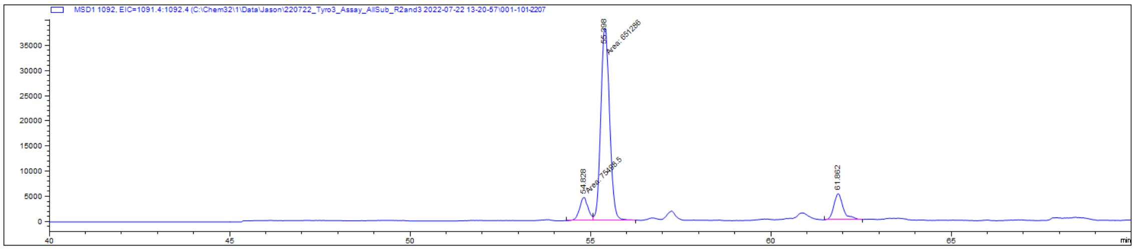

AXL Syn A at assay **t = 20 min; EIC of 1092**

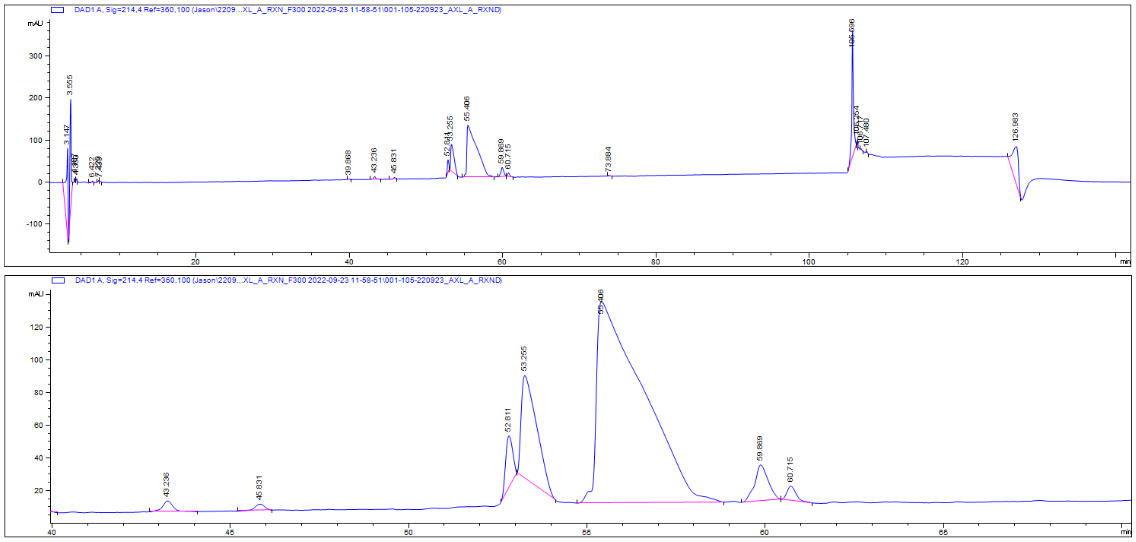

AXL Syn A TAM kinase reaction (upper); zoom of 214nm chromatogram (lower)

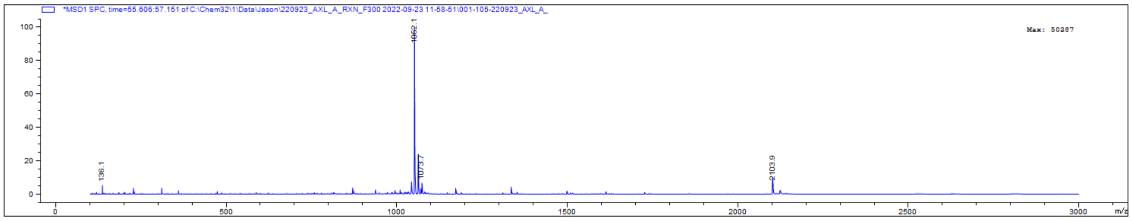

ID of peptide eluted at 55 min

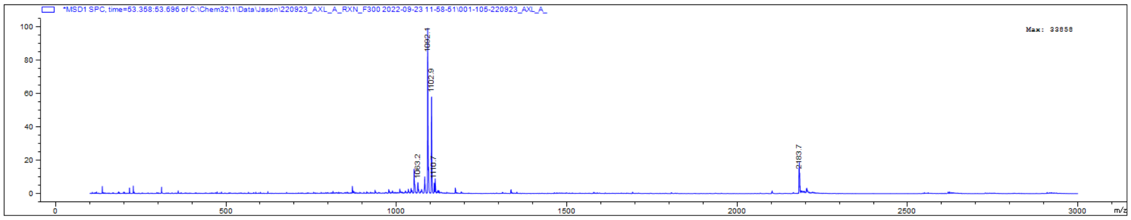

ID of peptide eluted at 53 min

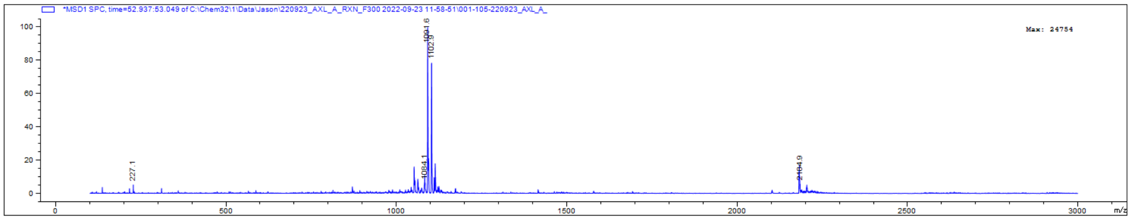

ID of peptide eluted at 52 min

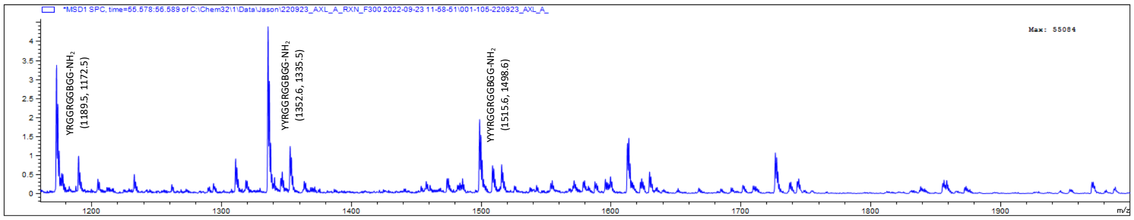

ID of peptide eluted at 55 min (zoom) = AXL Syn A (NDENNYYYRGGRGGBGG)

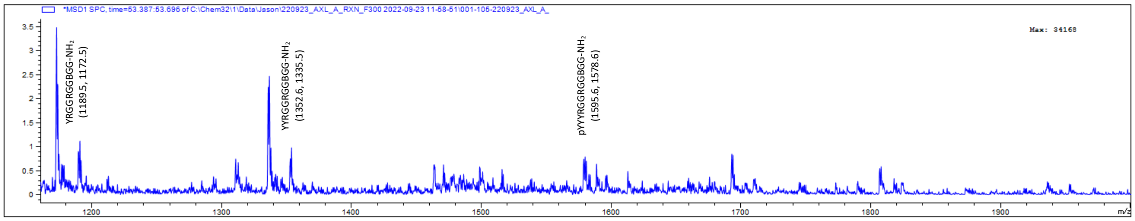


ID of peptide eluted at 53 min (zoom) = pAXL Syn A Major (NDENNpYYYRGGRGGBGG)

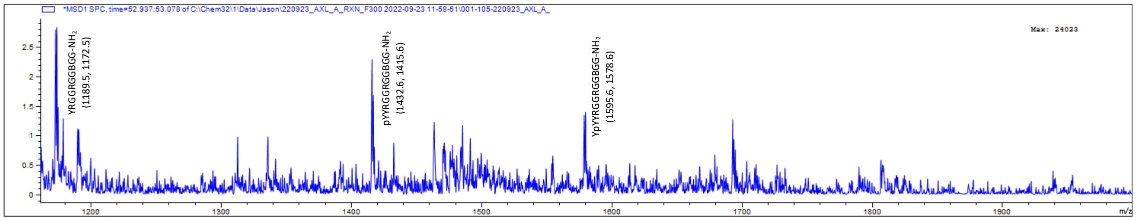


ID of peptide eluted at 52 min (zoom) = pAXL Syn A Minor (NDENNYpYYRGGRGGBGG)

#### *AXL Syn C:* NDENYYYYRGGRGGBGG

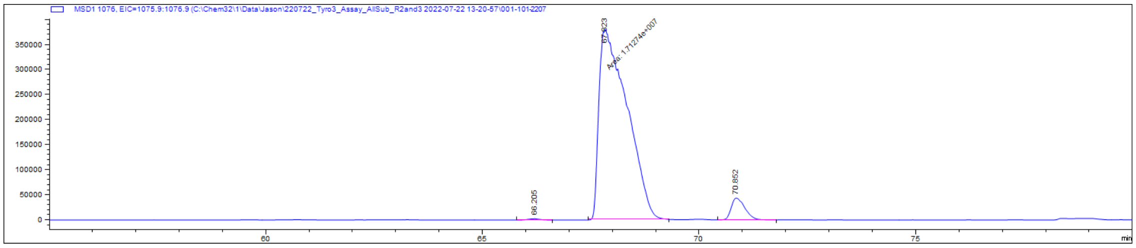


AXL Syn C at assay **t = 0 min; EIC of 1076**

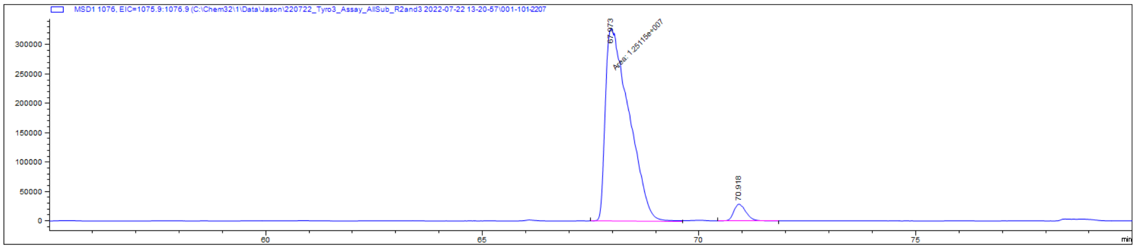


AXL Syn C at assay **t = 20 min; EIC of 1076**

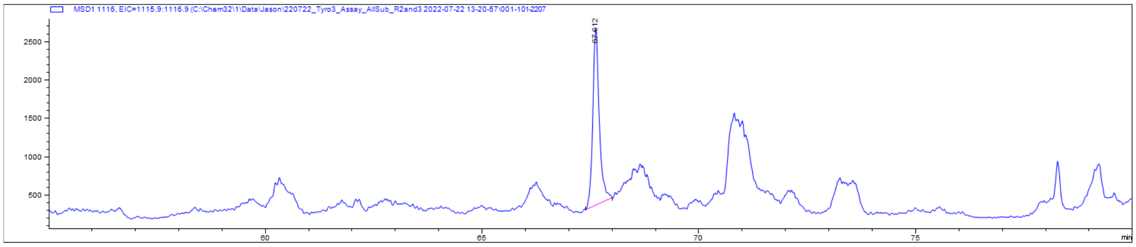

pAXL Syn C at assay **t = 0 min; EIC of 1116**

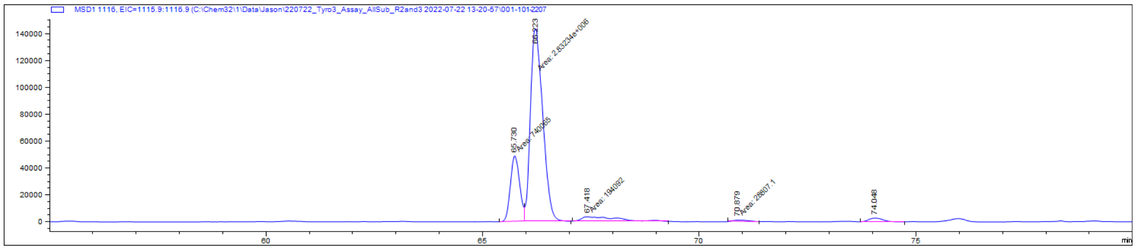

pAXL Syn C at assay **t = 20 min; EIC of 1116**

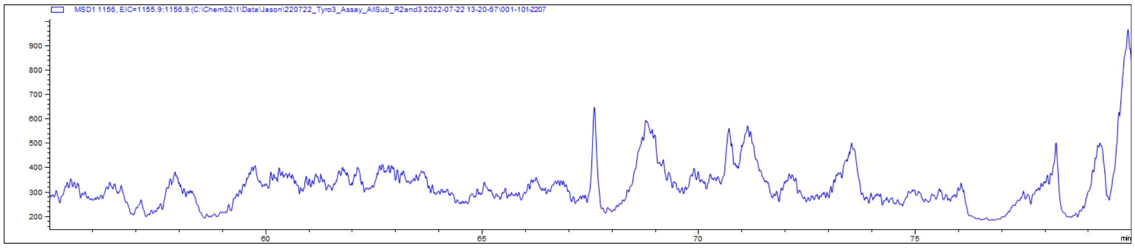

dpAXL Syn C at assay **t = 0 min; EIC of 1156**

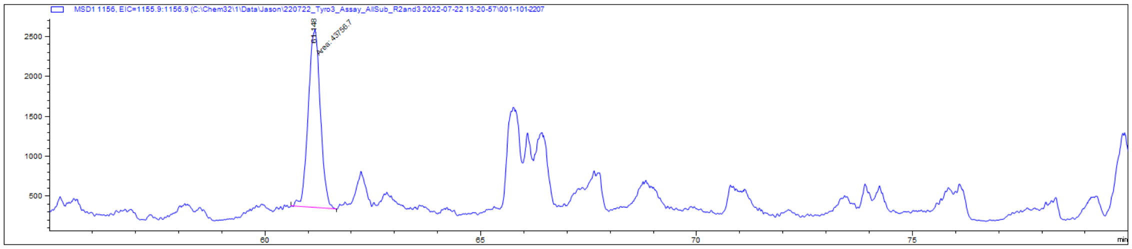

dpAXL Syn C at assay **t = 20 min; EIC of 1156**

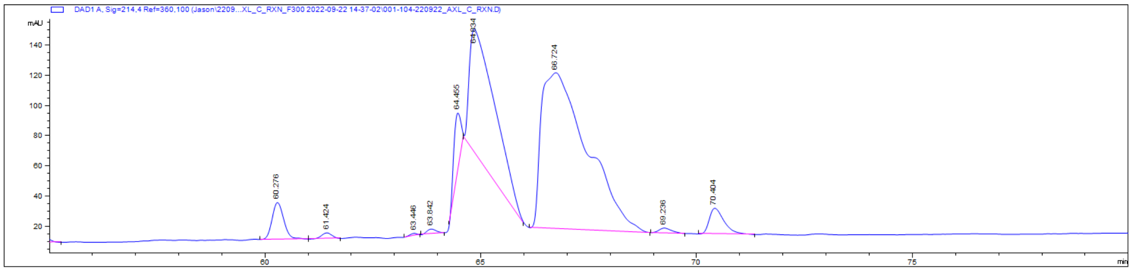

AXL Syn C TAM kinase reaction; zoom of 214nm chromatogram

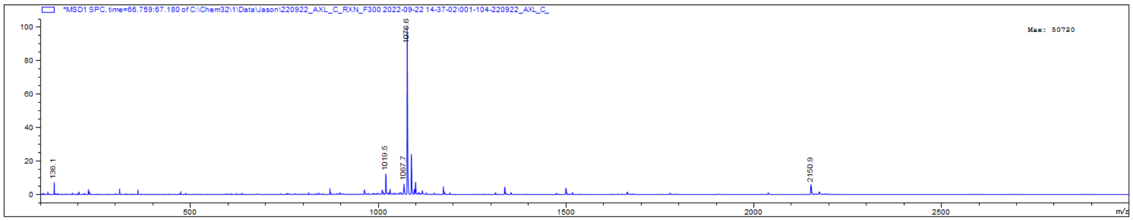

ID of peptide eluted at 67 min

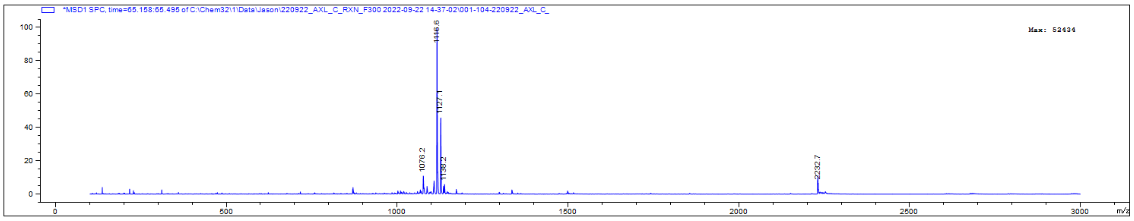


ID of peptide eluted at 65 min

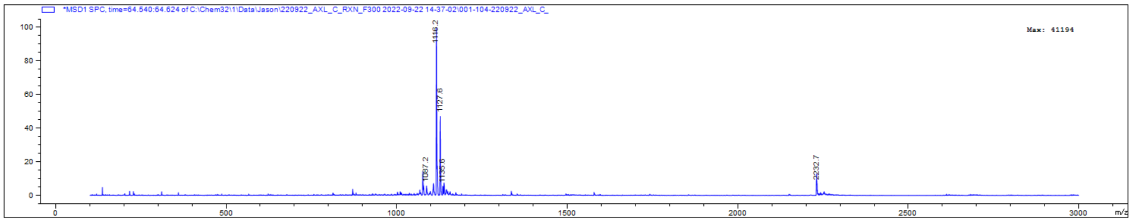


ID of peptide eluted at 64 min

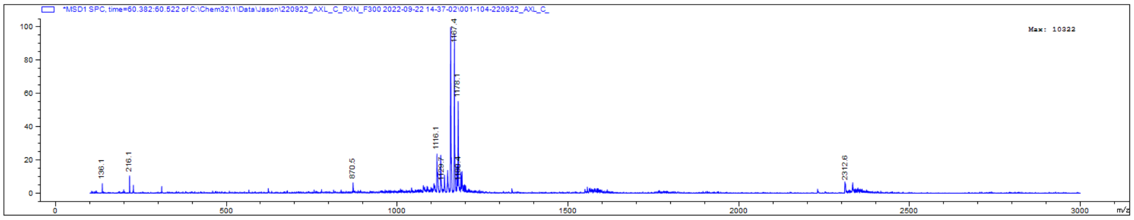


ID of peptide eluted at 60 min = dpAXL Syn C (NDENpYpYYYRGGRGGBGG)

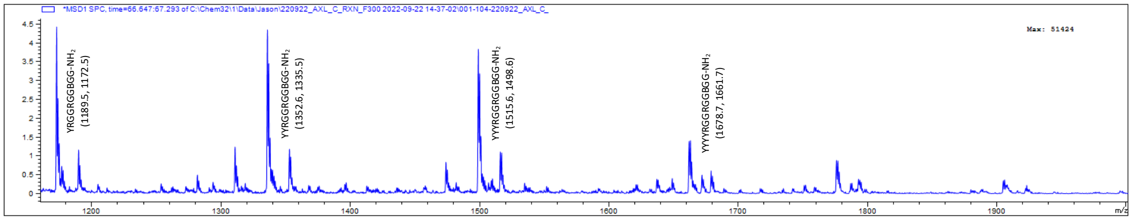


ID of peptide eluted at 67 min (zoom) = AXL Syn C (NDENYYYYRGGRGGBGG)

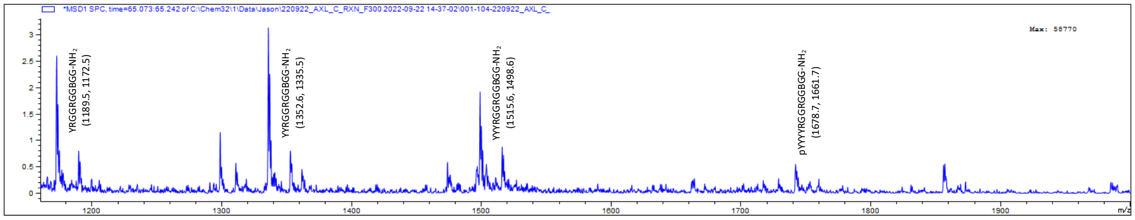


ID of peptide eluted at 65 min (zoom) = pAXL Syn C Major (NDENpYYYYRGGRGGBGG)

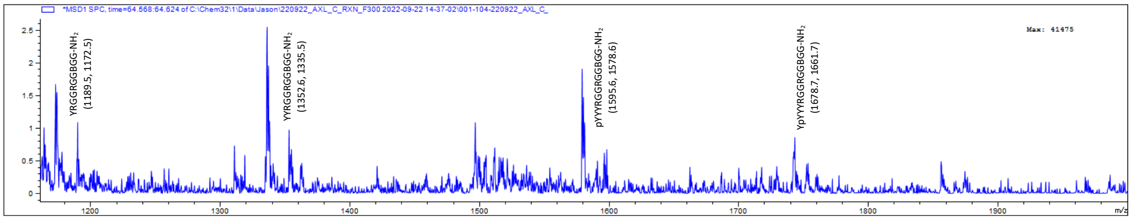


ID of peptide eluted at 64 min (zoom) = pAXL Syn C Minor (NDENYpYYYRGGRGGBGG)

*AXL Syn D:* NDENYYYYTGGRGGBGG

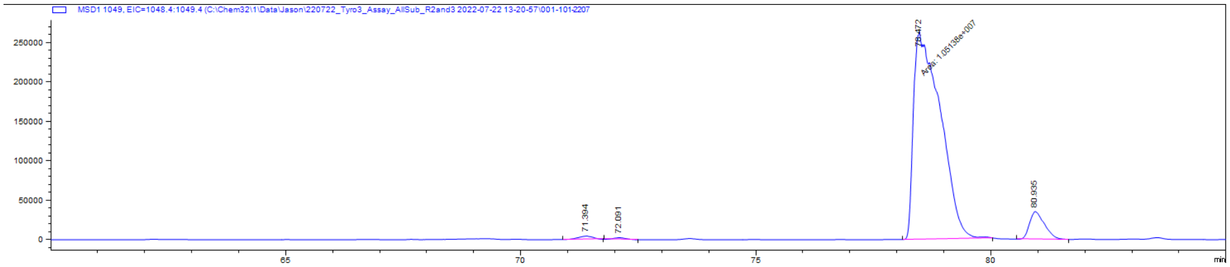

AXL Syn D at assay **t = 0 min; EIC of 1049**

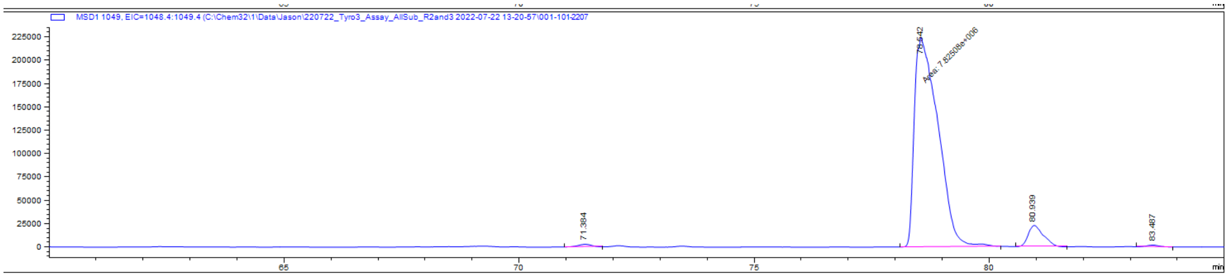

AXL Syn D at assay **t = 20 min; EIC of 1049**

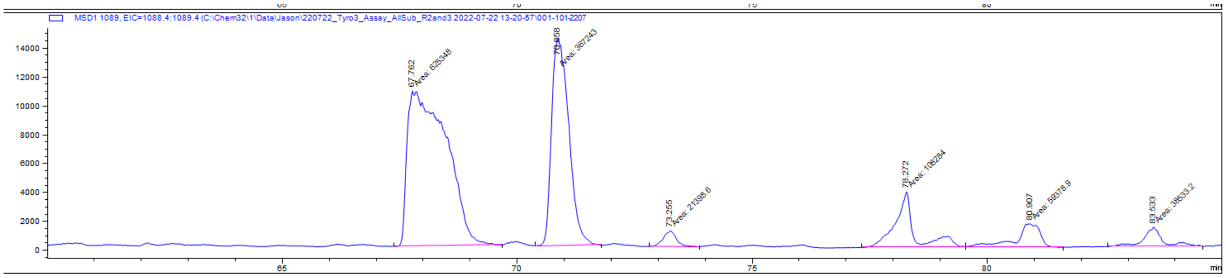

pAXL Syn D at assay **t = 0 min; EIC of 1089**

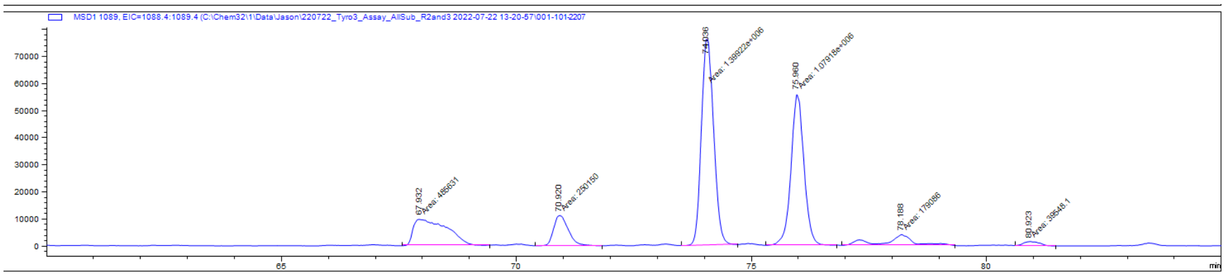

pAXL Syn D at assay **t = 20 min; EIC of 1089**

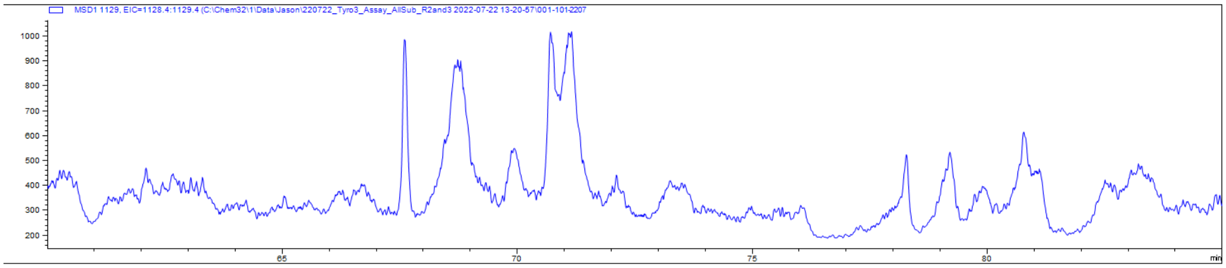


dpAXL Syn D at assay **t = 0 min; EIC of 1129**

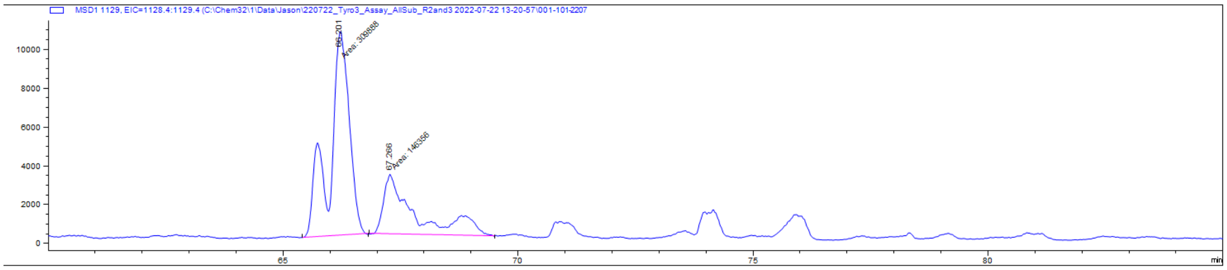


dpAXL Syn D at assay **t = 20 min; EIC of 1129**

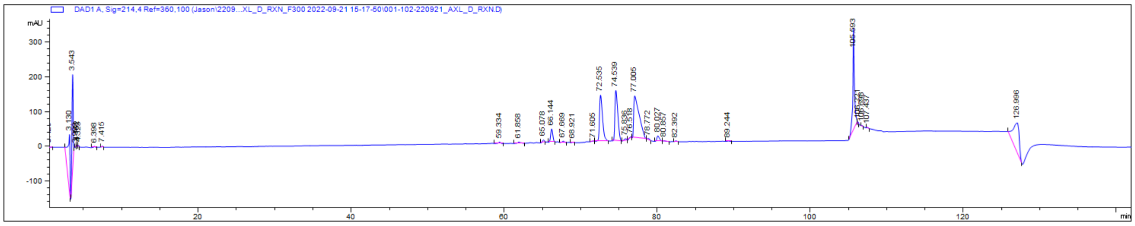


AXL Syn D TAM kinase reaction; full 214nm chromatogram

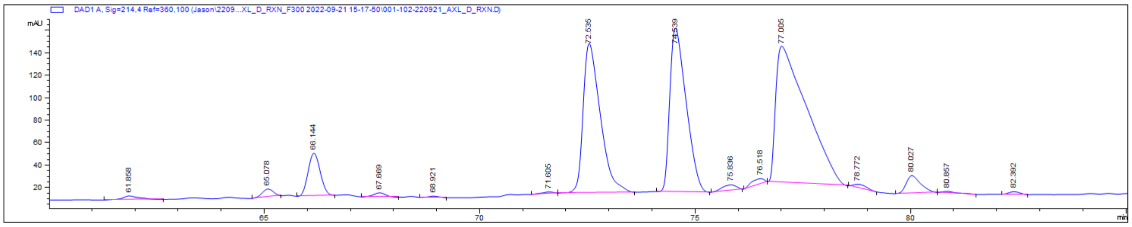


AXL Syn D TAM kinase reaction; zoom of 214nm chromatogram

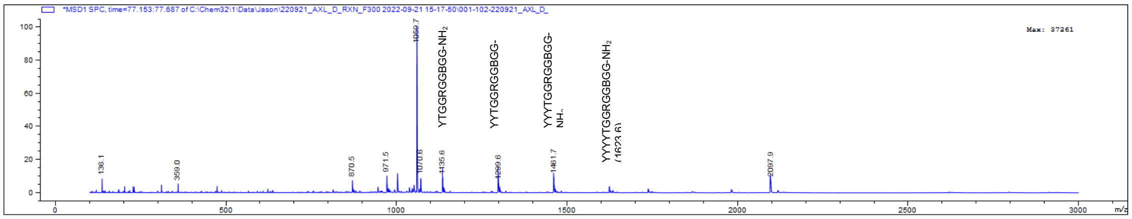


ID of peptide eluted at 77 min = AXL Syn D (NDENYYYYTGGRGGBGG)

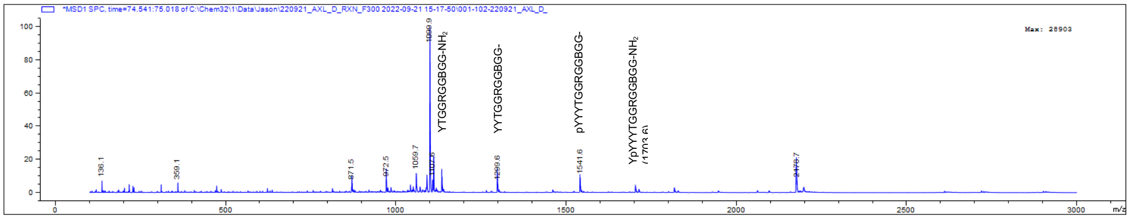


ID of peptide eluted at 74 min = pAXL Syn D2 (NDENYpYYYTGGRGGBGG)

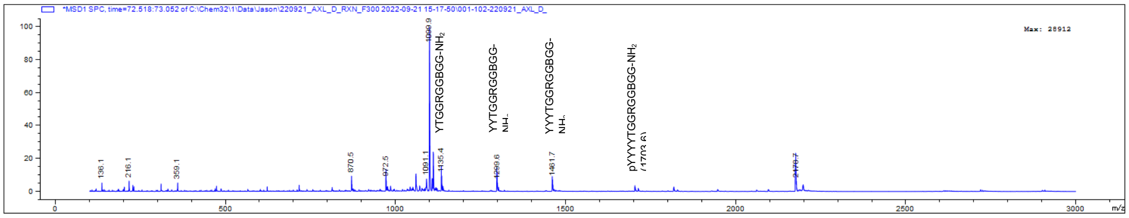


ID of peptide eluted at 72 min = pAXL Syn D1 (NDENpYYYYTGGRGGBGG)

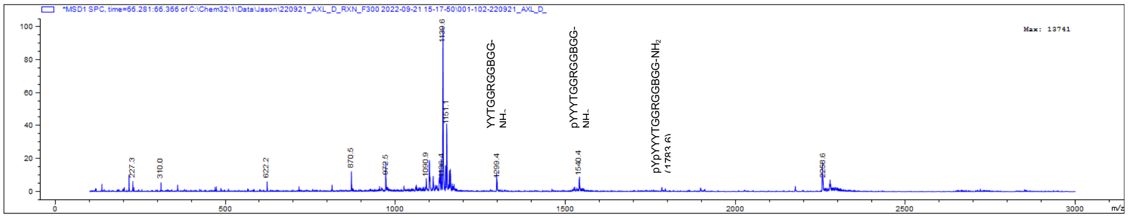


ID of peptide eluted at 66 min = dpAXL Syn D (NDENpYpYYYTGGRGGBGG)
